# mRNA localization is linked to translation regulation in the *Caenorhabditis elegans* germ lineage

**DOI:** 10.1101/2020.01.09.900498

**Authors:** Dylan M. Parker, Lindsay P. Winkenbach, Samuel P. Boyson, Matthew N. Saxton, Camryn Daidone, Zainab A. Al-Mazaydeh, Marc T. Nishimura, Florian Mueller, Erin Osborne Nishimura

## Abstract

*Caenorhabditis elegans* early embryos generate cell-specific transcriptomes despite lacking active transcription. This presents an opportunity to study mechanisms of post-transcriptional regulatory control. In seeking the mechanisms behind this patterning, we discovered that some cell-specific mRNAs accumulate non-homogenously within cells, localizing to membranes, P granules (associated with progenitor germ cells in the P lineage), and P-bodies (associated with RNA processing). Transcripts differed in their dependence on 3’UTRs and RNA Binding Proteins, suggesting diverse regulatory mechanisms. Notably, we found strong but imperfect correlations between low translational status and P granule localization within the progenitor germ lineage. By uncoupling these, we untangled a long-standing question: Are mRNAs directed to P granules for translational repression or do they accumulate there as a downstream step? We found translational repression preceded P granule localization and could occur independent of it. Further, disruption of translation was sufficient to send homogenously distributed mRNAs to P granules. Overall, we show transcripts important for germline development are directed to P granules by translational repression, and this, in turn, directs their accumulation in the progenitor germ lineage where their repression can ultimately be relieved.

**Summary:** Maternally loaded mRNAs localize non-homogeneously within *C. elegans* early embryos correlating with their translational status and lineage-specific fates.

## Introduction

The progression of life from two parental gametes into a developing embryo involves the transfer of gene expression responsibilities from the parental to zygotic genomes. In animals, a feature of this maternal-to-zygotic transition is a pause in transcription in late oogenesis, through fertilization, and during the first stages of zygotic development. Only after repression is relieved can the zygote’s genome autonomously transcribe [Hamm and Harrison, 2018, Robertson and Lin, 2015, Schulz and Harrison, 2019, Vastenhouw et al., 2019]. At this point, cell-specific promoter activity can diversify cell-specific transcriptomes through differential *de novo* production of mRNAs. However, before that point, any cell-specific differences in transcriptomes only arise through post-transcriptional mechanisms acting on mRNAs inherited from the parental gametes.

In *Caenorhabditis elegans*, transcriptional repression initiates in late oogenesis by an unknown mechanism [Gibert et al., 1984, Walker et al., 2007], but is sustained in post-fertilization stages by sequestration of transcriptional machinery to the cytoplasm [Guven-Ozkan et al., 2010]. Transcription resumes two-hours post-fertilization, initiating in the somatic cells of 4-cell embryos and culminating in the P_4_ cell of the primordial germ lineage (P lineage) at the 28-cell stage [Seydoux and Fire, 1994, Seydoux et al., 1996]. The P lineage is last to initiate transcription as it maintains transcriptional repression longer than somatic cells through inhibition of RNA Pol II initiation and elongation [Seydoux et al., 1996, Batchelder et al., 1999, Zhang et al., 2003, Ghosh and Seydoux, 2008, Guven-Ozkan et al., 2010] and reviewed in [Robertson and Lin, 2015].

Even in the absence of *de novo* zygotic transcription, the transcriptomes of early *C. elegans* blastomeres diversify. Single-cell resolution RNA-seq assays have determined that after the initial mitotic event, the two daughter cells (AB and P_1_) contain 80 AB-enriched and 201 P_1_-enriched transcripts distinguishing them [Osborne Nishimura et al., 2015]. Single-cell transcriptome profiles of blastomeres through the first four cell divisions identified additional maternally inherited transcripts with biased representation in different lineages [Tintori et al., 2016]. These cell-specific transcripts likely arise through post-transcriptional mechanisms such as differential rates of mRNA decay, mRNA stabilization, or by movement (active or passive) of transcripts into distinct regions of dividing cells.

Interestingly, there is no reason *a priori* for transcriptome diversification to be required for cell-specific protein production. Translational control is thought to play the major role in driving protein production during germline development [Merritt et al., 2008] and into early embryogenesis. Indeed, a major class of mutants that affect early cell fate development are cell-specific RNA Binding Proteins (RBPs), whose target transcripts are translated with spatiotemporal specificity [D’Agostino et al., 2006, Jadhav et al., 2008, Oldenbroek et al., 2012, Oldenbroek et al., 2013].

Still, the mRNA that encodes NEG-1 (Negative Effect on Gut development, a cell fate determinant) has an anterior bias preceding anterior NEG-1 protein production, suggesting that biases in mRNA can precede or even be amplified at the translation step [Elewa et al., 2015, Osborne Nishimura et al., 2015]. For these reasons, maternal mRNAs showing asymmetric representation in early embryonic cells may represent a class of transcripts that undergo post-transcriptional regulation important for cellular diversification. In this study, we explore the mechanisms and functions of this patterning.

We report that several maternally inherited transcripts localize to subcellular regions within individual cells. In general, the anterior-biased (AB cell-enriched) transcripts we surveyed tended to localize to cell-peripheral regions, often where the proteins they encode function. In contrast, posterior-biased (P_1_ cell-enriched) transcripts formed clustered granules overlapping with P granules, membrane-less compartments of RNAs and proteins that form liquid-liquid phase separated condensates or hydrogels (recently reviewed in [Seydoux, 2018, Marnik and Updike, 2019]).

Understanding the roles of P granules (and other phase-separated condensates)) in localizing mRNAs and impacting their expression is a major challenge in the field. It is possible that transcripts, like *nos-2* and others, associate with P granules for the purpose of forcing their translational repression. What is known is that worms can recover from P granule disruption in early embryonic stages to properly specify the germline [Gallo et al., 2010], but dysregulation of P granules that continues or occurs at later stages leads to perturbations in germ cell development [Wang et al., 2014], disruption of gene expression regulatory control [Campbell and Updike, 2015, Updike et al., 2014, Voronina et al., 2012] and fertility defects [Kawasaki et al., 2004, Spike et al., 2014, Wang et al., 2014]. In early embryos, P granules are large, cytoplasmic, and highly dynamic [Hird et al., 1996, Strome and Wood, 1982], but later grow into germ granules that coalesce around the nucleus [Sheth et al., 2010] forming an extended nuclear pore complex environment and branching into other microenvironment condensates such as mutator foci [Phillips et al., 2012] and Z-granules [Wan et al., 2018].

P granules house a growing number of identified protein components, but less is known about the RNA molecules within them. *cey-2*, *gld-1*, *mex-1*, *nos-2*, and *pos-1* mRNAs occupy P granules during embryonic stages [Jud et al., 2007, Subramaniam and Seydoux, 1999]. *gld-1*, *mex-1*, *nos-2*, *par-3*, *pos-1*, and *skn-1* mRNAs occupy germ granules in adults, and to a greater extent, in arrested oocytes where granules become large [Schisa et al., 2001, Subramaniam and Seydoux, 1999]. In general, germ granules are enriched in oligo-dT and SL1-splice leader mRNAs and depleted (or at least unconcentrated) for rRNA [Schisa et al., 2001]. By expanding the list of P granule-associated transcripts, we hope to use their identities and properties to better understand the link between P granules, mRNA localization, and regulatory control.

Many of the P granule-associated transcripts we identified were undergoing low or decreasing levels of translation. Indeed, the well-studied, P granule-resident mRNA *nos-2* is translationally repressed at early embryonic stages [D’Agostino et al., 2006, Jadhav et al., 2008, Subramaniam and Seydoux, 1999]. It is possible that mRNA transcripts, like *nos-2* and others, associate with P granules to promote their translational repression. In contrast, it is also possible that transcripts with low translational activity accumulate in P granules as a downstream step. In this study, we find that translational repression of *nos-2* mRNA precedes *nos-2* mRNA accumulation in P granules and can persist without P granule localization, supporting the second model. Further, we found that forcing translational repression on homogenously distributed transcripts directs them to P granules, again suggesting localization is a downstream step.

Overall, our work expands the list of membrane-associated mRNAs from 0 to 5 and P granule-associated from roughly 10 to 16. Given our rate of discovering subcellular patterning among maternally inherited transcripts (as opposed to zygotic transcripts), it is likely that maternally inherited transcripts are enriched for subcellular patterning. By identifying and studying more numerous mRNAs with subcellular localization in the *C. elegans* early embryo, we can better determine mechanisms and purposes of their localization in early development.

## Results

### Maternally inherited mRNA transcripts display subcellular localization

Single-cell RNA-seq assays have identified transcripts that are differentially abundant between cells prior to the onset of zygotic transcription in *C. elegans* [Hashimshony et al., 2012, Hashimshony et al., 2015, Osborne Nishimura et al., 2015, Tintori et al., 2016]. To verify the cell-specificity of these mRNAs and visualize their localization, we selected several to image in fixed *C. elegans* embryos using single-molecule resolution imaging (smFISH or smiFISH). We chose 8 AB-enriched transcripts, 8 P_1_-enriched transcripts, 4 uniformly distributed (maternal) transcripts, and 8 zygotically expressed transcripts. single-molecule resolution imaging confirmed the cell-specific patterning predicted by RNA-seq for 7 out of 8 AB-enriched, 7 out of 8 P_1_-enriched transcripts, and 4 out of 4 symmetric transcripts. Strikingly, many maternally inherited transcripts yielded subcellular localization patterns in addition to cell-specific patterning (Table 1, Fig. 1, Fig. S1).

**Table 1.** A survey of early embryonic mRNA for localization patterns. We surveyed 20 maternally inherited mRNA for localization patterns by smFISH (and in some cases, smiFISH [Tsanov et al., 2016]). Eight transcripts identified as AB-enriched (blue), eight P_1_-enriched (green), and four symmetrically distributed (orange) in single-cell resolution RNA-seq data at the 2-cell stage were surveyed [Osborne Nishimura et al., 2015]. Eight zygotically expressed transcripts were also surveyed (grey) [Tintori et al., 2016]. As a control for P granule localization, *nos-2* mRNA was included (yellow) [Schisa et al., 2001, Subramaniam and Seydoux, 1999].

| mRNA | maternal<br>v. zygotic | 2-cell enrichment by<br>RNA-seq (ranking) | 2-cell enrichment<br>by smFISH | patterning at 1-cell<br>to 16-cell by smFISH | notes |
| --- | --- | --- | --- | --- | --- |
| *erm-1* | maternal | AB-enriched (1) | AB-enriched | cell periphery |  |
| C50E13.3 | maternal | AB-enriched (3) | AB-enriched | no |  |
| *neg-1* | maternal | AB-enriched (4) | AB-enriched | no |  |
| *lem-3* | maternal | AB-enriched (7) | AB-enriched | cell periphery |  |
| *era-1* | maternal | AB-enriched (10) | AB-enriched | no |  |
| *ape-1* | maternal | AB-enriched (26) | symmetric | cell periphery |  |
| *mex-3* | maternal | AB-enriched (42) | AB-enriched | granular | granules are in the P-lineage |
| *tes-1* | maternal | AB-enriched (75) | AB-enriched | cell periphery | variable |
| *chs-1* | maternal | P<sub>1</sub>-enriched (1) | P<sub>1</sub>-enriched | granular |  |
| *clu-1* | maternal | P<sub>1</sub>-enriched (4) | P<sub>1</sub>-enriched | granular |  |
| *ipgm-1* | maternal | P<sub>1</sub>-enriched (25) | P<sub>1</sub>-enriched | granular | also known as F57B10.3 |
| *cpg-2* | maternal | P<sub>1</sub>-enriched (30) | P<sub>1</sub>-enriched | granular |  |
| *pgl-3* | maternal | P<sub>1</sub>-enriched (32) | P<sub>1</sub>-enriched | no |  |
| T24D1.3 | maternal | P<sub>1</sub>-enriched (40) | P<sub>1</sub>-enriched | granular |  |
| *puf-3* | maternal | P<sub>1</sub>-enriched (75) | symmetric | granular |  |
| *bpl-1* | maternal | P<sub>1</sub>-enriched (170) | P<sub>1</sub>-enriched | no |  |
| *set-3* | maternal | symmetric | symmetric | no | granular in posterior cells at later stages |
| *gpd-2* | maternal | symmetric | symmetric | no |  |
| B0495.7 | maternal | symmetric | symmetric | no |  |
| *imb-2* | maternal | symmetric | symmetric | no |  |
| *elt-2* | zygotic |  |  | no |  |
| *end-1* | zygotic |  |  | no |  |
| *hlh-27* | zygotic |  |  | no |  |
| *hsp-60* | zygotic |  |  | no |  |
| *ref-1* | zygotic |  |  | no |  |
| *tbx-32* | zygotic |  |  | no |  |
| *tbx-38* | zygotic |  |  | no |  |
| Y75B12A.2 | zygotic |  |  | no |  |
| *nos-2* | maternal | symmetric | symmetric | granular | previously reported P granule mRNA |

**Figure 1.**
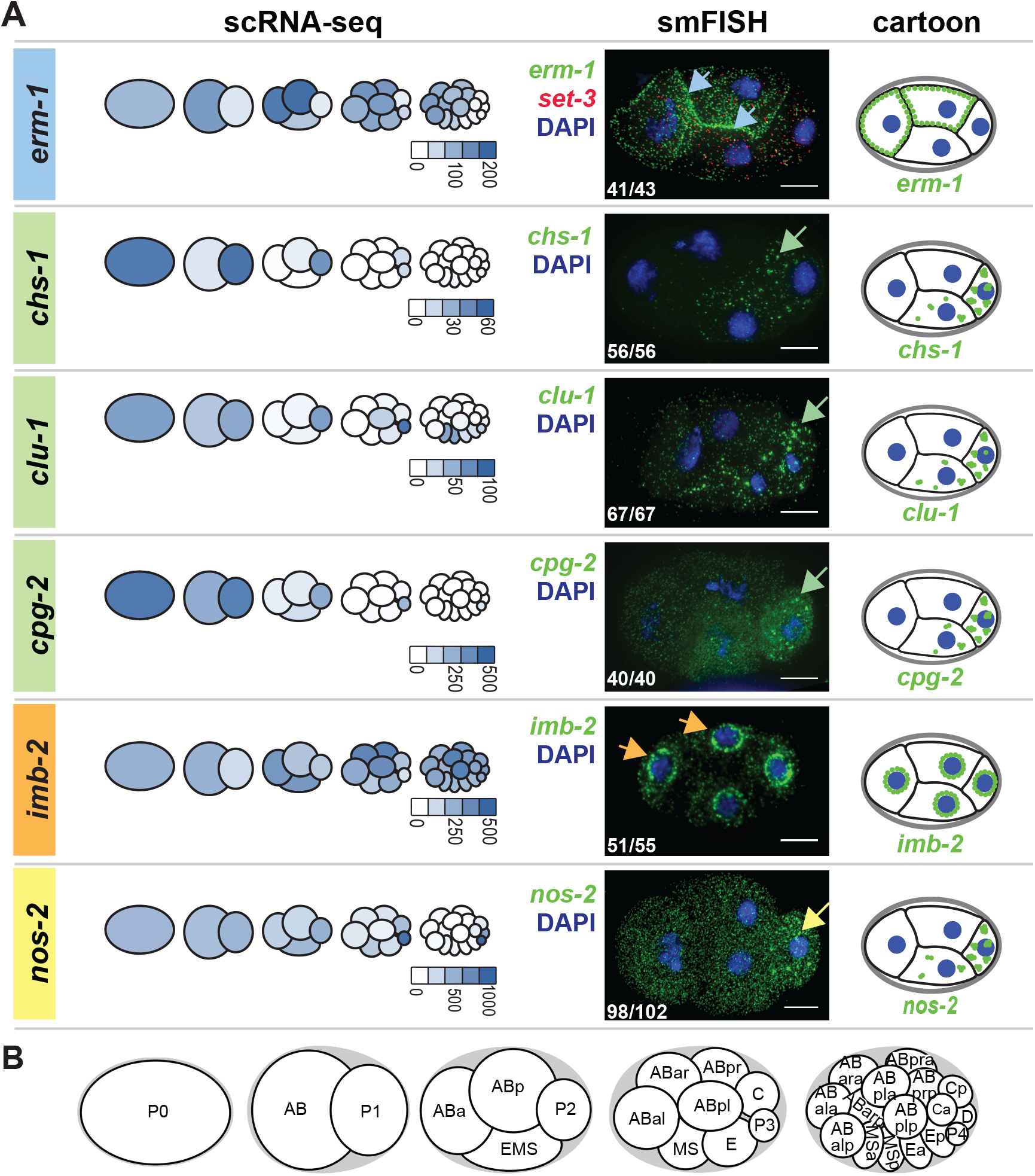
Subcellular localization patterns of maternally inherited mRNAs. **(A)** mRNA localization patterns for *erm-1, chs-1, clu-1, cpg-2, imb-2*, and *nos-2* are shown (Table 1, Fig. S1). They represent AB-enriched (blue), P_1_-enriched (green), and symmetric (orange) maternal mRNAs and a known P granule control (yellow). **Left:** mRNA abundance through the first four cell divisions as previously reported by single-cell RNA-seq data [Tintori et al., 2016] is illustrated as pictographs with normalized transcript abundance values indicated to the lower right of each pictograph. **Center:** mRNA were imaged by smFISH, and a representative 4-cell stage image for each shows the transcript of interest (green), DNA (DAPI, blue). *set-3* (red) was co-probed in each embryo, but is only shown in one for simplicity. mRNAs were concentrated at cell peripheries (*erm-1*, blue arrows), clusters (*chs-1, clu-1*, and *cpg-2*, green arrows), nuclear peripheries (*imb-2*, orange arrows), or at known P granules (*nos-2*, yellow arrow). Inset numbers represent the number of times patterning was observed out of the total 4-cell stage embryos surveyed. Scale bars represent 10 *μ*m. **Right:** Cartoon depictions of each mRNA of interest (green) are shown to emphasize subcellular distribution patterns. **(B)** Cartoon depictions of the first five embryonic stages.

AB-enriched transcripts tended to localize to cell peripheries (Table 1). Specifically, AB-enriched *erm-1 (Ezrin/Radixin/Moesin)*, *lem-3 (LEM domain protein)*, *ape-1 (APoptosis Enhancer)*, and *tes-1 (TEStin homolog)* mRNAs accumulated there. ERM-1 protein also accumulates at cell-to-cell contacts where it functions in the remodeling of apical junctions [Van Fürden et al., 2004]. Similarly, LEM-3, a nucleic acid metabolizing enzyme, localizes to cell membranes (supplemental material of [Dittrich et al., 2012]) and cytoplasmic foci. The localization of APE-1 and TES-1 proteins are uncharacterized, but they contain domains known to associate with membranes (ankyrin-repeat domain in APE-1 and PET domain in TES-1) [Bennett and Baines, 2001, Sweede et al., 2008]. For this paper, we focused on *erm-1* as a representative of this group (Fig. 1).

P_1_-enriched transcripts tended to aggregate into RNA granules in the P lineage (Table 1, Fig. 1 Fig. S1). This included transcripts important in eggshell formation such as *chs-1 (CHitin Synthase)* and *cpg-2 (Chondroitin ProteoGlycan)*, mitochondrial distribution and stress response such as *clu-1 (yeast CLU-1 [CLUstered mitochondria] related)*, as well as the carbohydrate metabolizing enzyme *ipgm-1* (*cofactor-Independent PhosphoGlycerate Mutase homolog*) (recently renamed from F57B10.3) [Fields et al., 1998, Maruyama et al., 2007, Olson et al., 2012, Zhang et al., 2004].

Of the maternally inherited transcripts that distribute symmetrically at the 2-cell stage, only one of the four we tested showed subcellular patterning (Table 1, Fig. S1). The transcript *imb-2 (IMportin Beta family)* localized to nuclear peripheries, coincident with the location of its encoded protein, an Importin-*β* homologue that facilitates nuclear pore complex import (Fig. 1). In no cases did we observe subcellular localization for mRNAs expressed zygotically suggesting that maternally loaded mRNAs may be over-represented for subcellular localization (Table 1).

In addition to these surveyed transcripts, we also used smFISH to image *nos-2 (NanOS related)*, a previously reported mRNA resident of P granules required for germline maintenance and fertility [Subramaniam and Seydoux, 1999] (Table 1, Fig. 1). As expected, smFISH verified P granule localization of *nos-2* mRNA and showed granular patterning was coincident with P lineage enrichment – both beginning at late 4-cell stage (Fig. 2).

**Figure 2.**
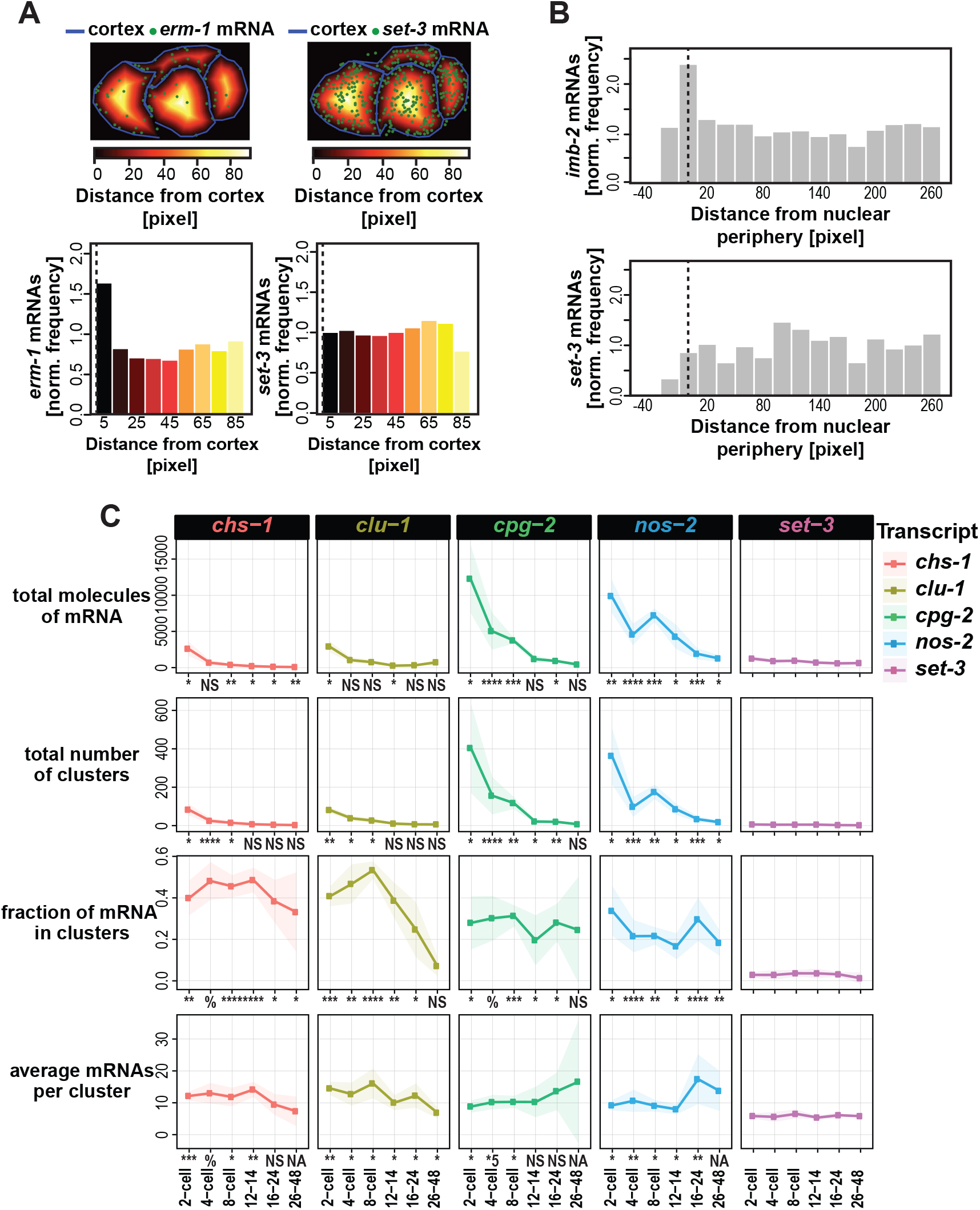
Quantification of mRNA and their patterning. **(A)** The number of mRNA molecules (green dots) located within binned distances from the cell cortex (blue lines) were tabulated and normalized to the volume of each concentric space. The frequencies with which *erm-1* mRNA and *set-3* mRNA occurred at varying distances are shown. **(B)** The frequencies with which mRNA appeared in relation to the nuclear peripheries were similarly calculated for *imb-2* mRNA and *set-3* mRNA. **(C)** Several metrics of clustering were quantified for: *chs-1* (red), *clu-1* (ochre), *cpg-2* (green), the known P granule resident *nos-2* (blue), and the symmetric comparison *set-3* (purple). We calculated the 1) total number of RNAs in each embryo, 2) total number of clusters identified in each embryo, 3) fraction of total mRNAs within clusters, and 4) the average estimated number of mRNA molecules per cluster within each embryo. The average of each metric and their standard deviation (shading) for each transcript at five cell stages are shown representing a minimum of 5 embryos assayed. Significance indicates p-values derived from multiple test corrected t-tests (NS > 0.05; 0.05 > * > 0.005; 0.005 > ** > 0.0005; 0.0005 > *** > 0.00005; 0.00005 > **** > 0.000005; 0.000005 > *5 > 0.0000005; >%< 0.0000005)

To explore the dynamics of subcellular patterning through embryogenesis, we imaged key transcripts from the 1-cell stage through hatching. The onset and persistence of subcellular mRNA localization varied depending on the transcript and its biology (Fig. 2). *chs-1* mRNA first localized to posterior clusters at the 1-cell or 2-cell stage but degraded over successive cell divisions until eventually completely dissipating by the 48-cell stage (Fig. 2) whereas *imb-2* appeared at or near nuclear membranes in all stages assayed. This is consistent with the roles of the proteins as CHS-1 is essential primarily for deposition of chitin in the eggshell between oogenesis and egg-laying [Zhang et al., 2005] whereas the IMB-2 protein is required throughout the life of the worm for nuclear import [Putker et al., 2013]. In contrast to *chs-1*, *nos-2* mRNA distributed homogeneously prior to the 4-cell stage of development. In the late 4-cell stage, *nos-2* mRNA gathered into clusters in the P lineage coincident with its degradation in somatic cells. These *nos-2* mRNA clusters grew in size until the 28-cell stage (Fig. 2). At the 28-cell stage, *nos-2* transcripts became visible as individuals in the cytoplasm concurrent with a decrease in the size of *nos-2* mRNA clusters. Translational regulation of *nos-2* is dynamic during these stages. *nos-2* mRNA is translationally repressed prior to the 28-cell stage. That repression is relieved at the 28-cell stage [D’Agostino et al., 2006, Jadhav et al., 2008]. Therefore, the transition in RNA localization accompanies this transition in regulatory status. What was more surprising is that *nos-2* mRNA could both be observed free floating as individual mRNAs and localized into granules prior to the 28-cell stage during its phase of translational repression. During the 1-cell, 2-cell, and early 4-cell stages, *nos-2* mRNA fails to produce protein, but also does not localize to clusters, illustrating that these processes can be uncoupled. Altogether, subcellular transcript localization appears transient or persistent depending on the encoded function of the mRNA.

### Quantification strategies to characterize mRNA patterning

To better describe the subcellular mRNA patterns we observed, we detected individual mRNA molecules in 3D images using FISH-quant [Mueller et al., 2013] and developed metrics to describe their localizations at membranes or within clusters.

*erm-1* mRNA appeared to localize to cell peripheries. To characterize this propensity in an unbiased manner, we calculated the frequency with which *erm-1* transcripts accumulated at increasing distances from cell membranes (Fig. 2A). After normalizing for the decreasing volumes of each concentric space, we determined *erm-1* mRNA were twice as likely to occur within 5 microns of a cell membrane versus greater than 5 microns from one. In contrast, homogenously distributed *set-3* transcripts were equally likely to be present at all distances (both measured using 10-micron bin sizes) (Fig. 2A).

Similarly, we calculated the frequency of *imb-2* mRNA at increasing distances from the nuclear periphery (Fig. 2B). *imb-2* transcripts were twice as abundant within 10 microns from the nuclear membrane versus at 10 microns or more from a nuclear membrane, again adjusting for volumes of these spaces. The more ubiquitous *set-3* transcripts showed no nuclear peripheral-enrichment.

In developing metrics to describe features of mRNA clusters, we found that overlapping mRNA signals complicated the “single molecule” nature of smFISH which relies on sufficient spacing between individual transcripts. To overcome this, we used a tiered approach, first identifying individual mRNAs [Mueller et al., 2013] and secondly applying the fluorescence intensities and volumes of the individuals to fit a Gaussian Mixture Model (GMM) that estimates the number of molecules contributing to signal overlap (see Methods). Deconvolved mRNA molecules could then be separated into clusters using a geometric nearest neighbor approach [Ester, M., Kriegel, H. P., Sander, J., & Xu, 1996].

To characterize mRNA clusters, we quantified the 1) the total number of mRNA molecules per embryo, 2) the total number of mRNA clusters per embryo, 3) the fraction of total mRNAs that localize into clusters (as opposed to individuals), and 4) the estimated number of mRNAs within each cluster. We calculated these measurements for four clustered transcripts (*chs-1, clu-1, cpg-2*, and *nos-2*) at five stages of embryonic development (Fig. 2C). This revealed differences between transcripts. For example, *cpg-2* and *nos-2* were the most abundant (~10,000 molecules per embryo) in contrast to *chs-1* or *clu-1* (~2,500 molecules per embryo) at the same timepoint (2-cell stage). The number of *cpg-2* and *nos-2* mRNA molecules comprising each cluster increased over time whereas *chs-1* and *clu-1* did not. For *nos-2*, mRNA accumulated to a maximum of 20 molecules per cluster or greater at the 24-cell stage, just prior to *nos-2*’s translational activation. After this point, *nos-2* mRNA clusters decreased in size, appearing dispersed in the cytoplasm. All clustered transcripts exhibited marked differences in clustering statistics from the homogenously distributed *set-3* transcripts.

### Clustered transcripts *chs-1*, *clu-1*, *cpg-2* and *nos-2* colocalize with markers of P granules and, less frequently, with markers of P-bodies

mRNA clustering is typically indicative of localization into granules, membrane-less compartments that concentrate mRNAs and proteins into liquid-liquid phase-separated condensates or hydrogels. Many types of condensates exist, such as stress granules (associated with translationally repressed transcripts that accumulate during stress), P-bodies (Processing bodies, associated with RNA processing enzymes), and germ granules (associated with regulatory control in animal germ cells). In *C. elegans* germ granules are specifically called P granules in the early embryo (Fig. 3A) (recently reviewed in [Seydoux, 2018, Marnik and Updike, 2019]) where they differ from later-stage germ granules in that they are free floating in the cytoplasm and concentrate down the P lineage with each successive cell division. Dual mechanisms of preferential coalescence/segregation in the P lineage and disassembly/degradation in somatic cells drives their concentration in the P lineage [Brangwynne et al., 2009, DeRenzo et al., 2003, Wang et al., 2014].

**Figure 3.**
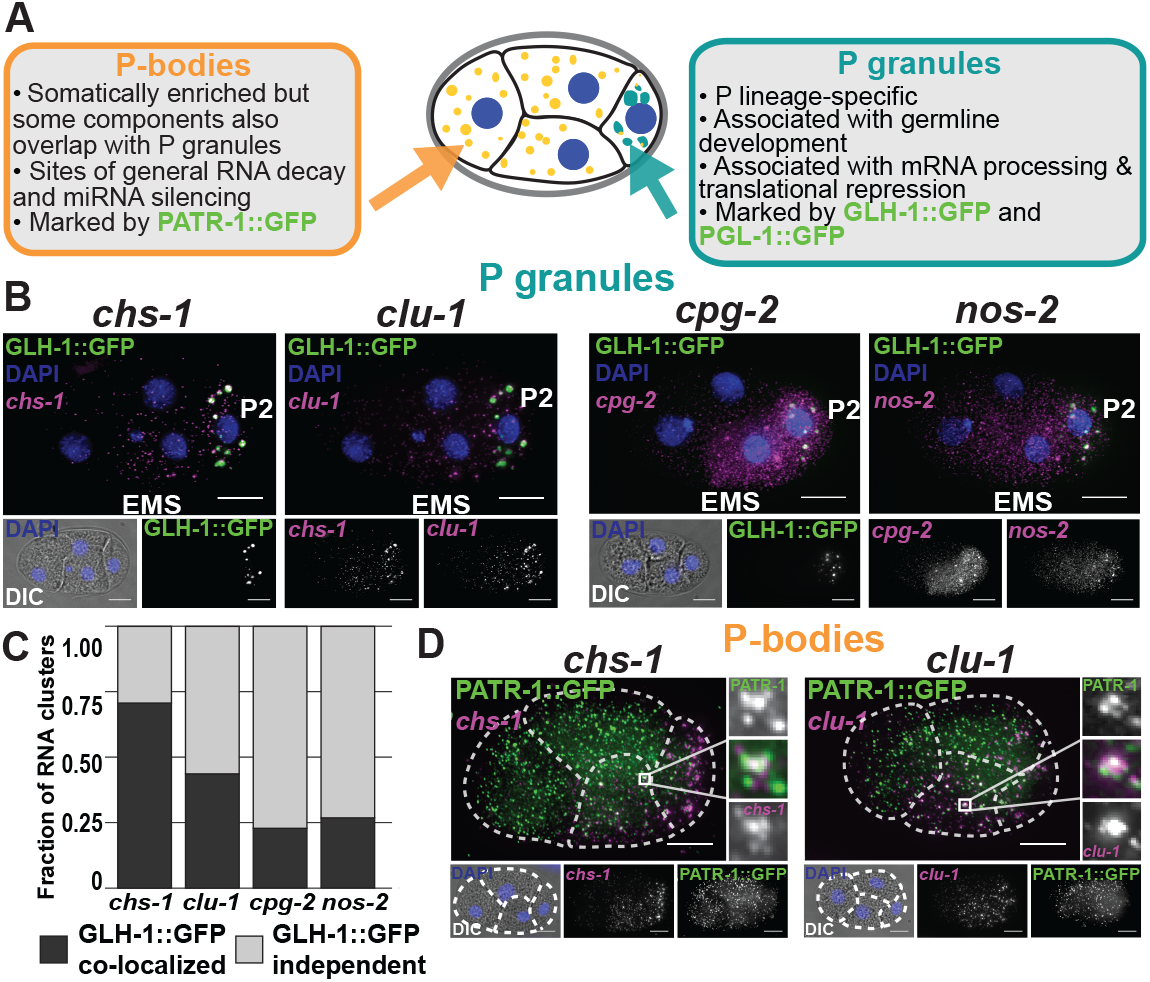
Posterior, clustered mRNAs co-localize with P granules and P-bodies. **(A)** P granules are distinct from P-bodies in their locations, functions, and marker proteins. **(B)** Fixed embryos were imaged for the P granule protein GLH-1∷GFP (green) and *chs-1, clu-1, cpg-2*, or *nos-2* transcripts (all in magenta). DNA (DAPI, blue) and DIC are also shown. **(C)** The fraction of mRNA clusters overlapping with P granules (dark grey) and P granule-independent clusters (light grey) was calculated by assessing spatial overlap between mRNA clusters and GLH-1∷GFP-marked P granules. **(D)** Fixed embryos were imaged for the P-body protein marker PATR-1∷GFP amplified using immunofluorescence (green) with smFISH imaging of *chs-1* or *clu-1* mRNA (magenta), and DNA (DAPI, blue). Outset images illustrate regions of co-localization.

Given that we observed *chs-1*, *clu-1*, and *cpg-2* mRNAs clustered and progressing down the P lineage, we hypothesized that they may be within P granules. To test this, we imaged *chs-1*, *clu-1*, *cpg-2*, and, for comparison, *nos-2* by smFISH in worms expressing P granule markers GLH-1∷GFP (Fig. 3B) or PGL-1∷GFP (Fig. 3). mRNA clusters overlapped with both P granule markers. 23% (*cpg-2*) to 75% (*chs-1*) of identified mRNA clusters overlapped with GLH-1∷GFP-marked P granules at the 4-cell stage, and their co-occurrence increased thereafter (Fig. 3C). Larger mRNA clusters were more likely to co-occupy space with P granules (Fig. 4). Conversely, 13 – 57% of GLH-1∷GFP marked P granules contained an mRNA cluster of any individually queried transcript, suggesting some heterogeneity in their content. Together, these findings illustrate that P-lineage enriched mRNA clusters in this study are P granule-associated RNAs.

**Figure 4.**
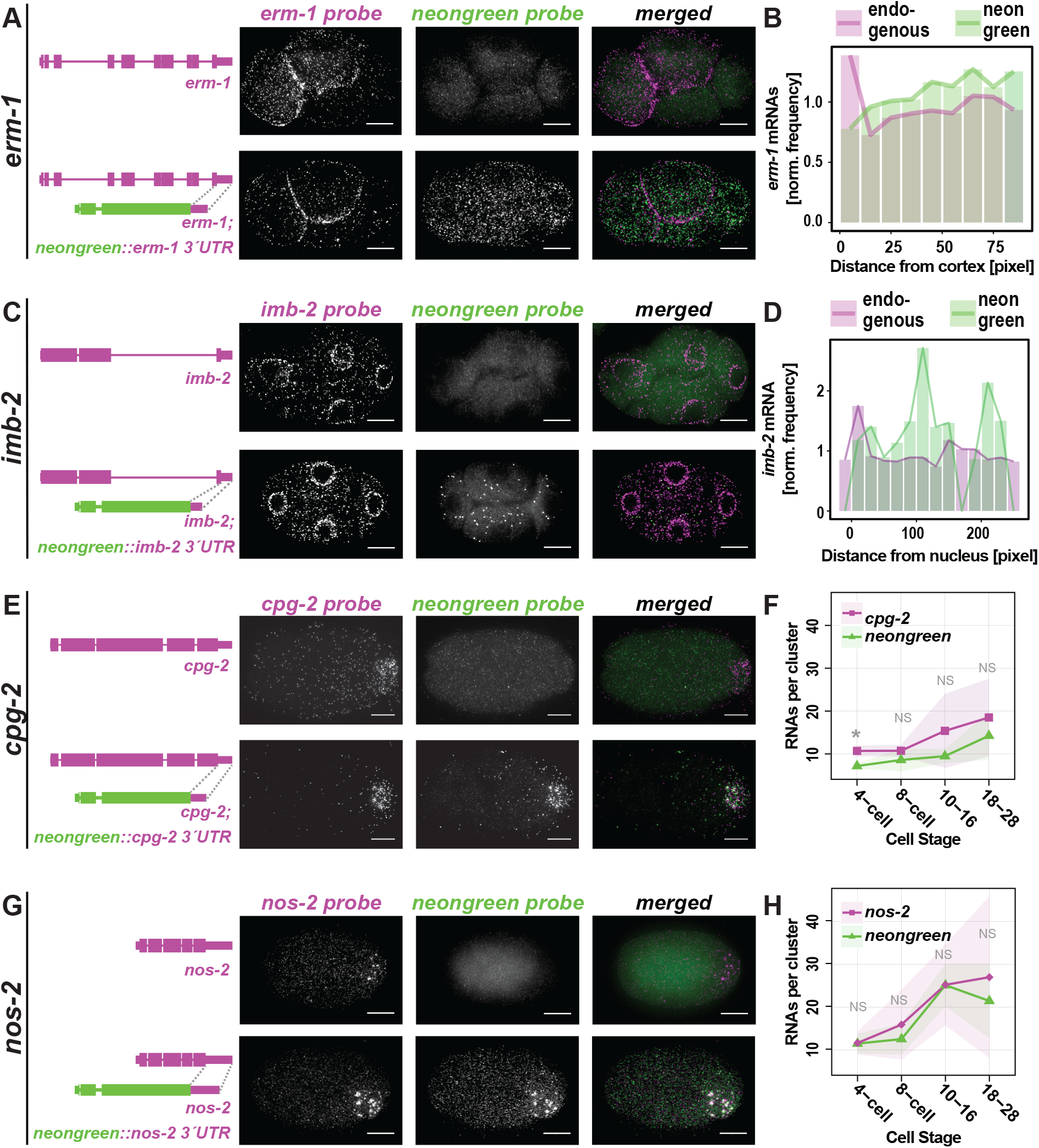
3’UTRs of clustered, but not membrane associated transcripts, are sufficient for subcellular localization. The 3’UTRs of **(A)** *erm-1*, **(B)** *imb-2*, **(E)** *cpg-2*, and **(G)** *nos-2* were appended to monomeric *NeonGreen (mex-5p∷mNeonGreen∷3’UTR of Interest)* and introduced as a single copy insert into otherwise wild-type worms to test whether 3’UTRs of interest were sufficient to drive subcellular localization patterns observed for full length transcripts. Wild-type control strains (top panels) and transgenic strains (bottom panels) were imaged by smFISH using probes hybridizing to the endogenous mRNA of interest (left), *mNeonGreen* mRNA (middle) and merged (right). Representative 4-cell stage embryos are shown with scale bars representing 10 *μ*m. **(B and D)** Quantification of images shown in (A and C) indicating the normalized frequency of (B) *erm-1*, or (D) *imb-2* mRNA and *mNeonGreen* mRNA at increasing distances from cell peripheries. (F and H) The estimated mRNA content per cluster from a minimum of 5 embryos at each of five binned stages of development are reported for endogenous (F) *cpg-2* or (H) *nos-2* (magenta) and *mNeonGreen* reporters (green). p-values from multiple test corrected t-tests are shown (NS > 0.05; 0.05 > * > 0.005)

Depending on the transcript, 25% to 75% of RNA clusters, typically smaller clusters, were distinct from P granule markers at the 4-cell stage. These occurred in P cells and their sisters (most evidently in EMS). Because many of the clustered mRNAs we are studying (*chs-1*, *clu-1*, *cpg-2*, and *nos-2*) degrade in early embryogenesis (Fig. 2C), we hypothesized that the RNA clusters that did not overlap with P granule markers were P-bodies. P-bodies, or Processing-bodies – as opposed to P granules – are associated with RNA decay as they contain high concentrations of RNA degrading proteins (DCAP-1, Argonaute, and Xrn-1) [Parker and Sheth, 2007] (Fig. 3A). In *C. elegans*, P granules and P-bodies share some protein components, but specific proteins distinguish each [Gallo et al., 2008, Voronina et al., 2011]. To test our hypothesis, we imaged *chs-1*, *clu-1*, *cpg-2*, and *nos-2* by smFISH concurrently with PATR-1∷GFP (yeast PAT-1 Related) amplified by immunofluorescence to mark P-bodies (See Materials and Methods, Fig. 5). *chs-1* and *clu-1* transcripts were enriched in posterior cells whereas PATR-1∷GFP predominantly localized to somatic cells. However, within their regions of overlap, we identified co-localized clusters indicating that some clusters of *chs-1* and *clu-1* mRNAs may reside within P-bodies (Fig. 3D). Many *chs-1* and *clu-1* mRNA clusters overlapped with neither P granule nor P-body markers, leaving their identity a mystery. Whether these mRNA clusters are stable or short-lived is currently unclear as fixed smFISH assays cannot resolve their dynamics.

**Figure 5.**
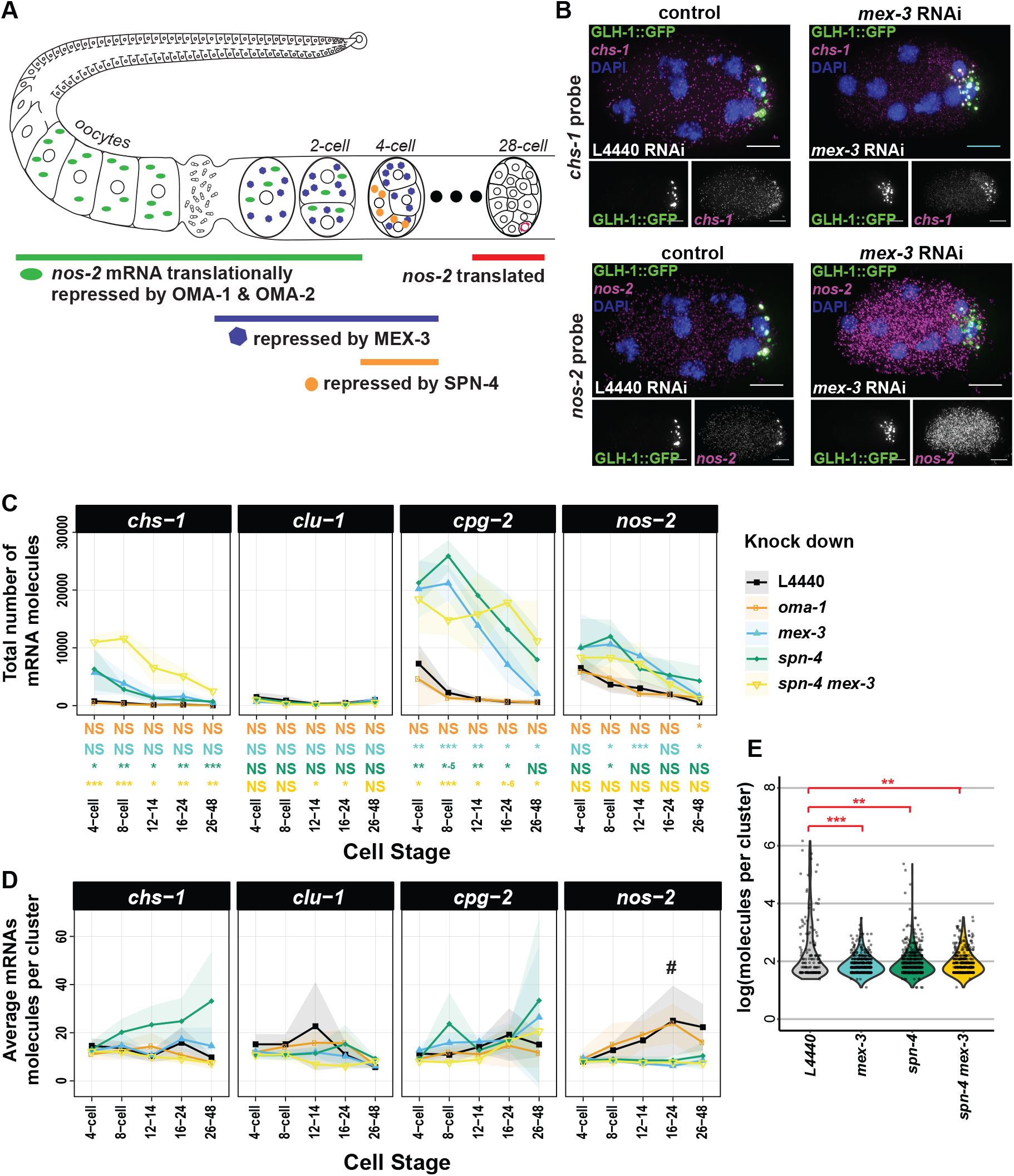
RNA Binding Proteins (RBPs) that repress translation of *nos-2* mRNA also impact degradation and subcellular localization of other mRNAs. **(A)** A succession of RBPs cooperatively repress *nos-2* translation from oogenesis through the 28-cell stage **(B)** Depletion of these RBPs impacted RNA abundance and localization of *nos-2* and other posterior, clustered mRNA transcripts. The impact of depleting the MEX-3 RBP by RNAi is shown for *chs-1* mRNA (magenta, top) and *nos-2* mRNA (magenta, bottom) as imaged by smFISH in a P granule marker strain (GLH-1∷GFP, green). **(C, D)** The (C) total number of mRNA molecules and (D) average number of mRNA molecules per cluster for four different RBP knockdown conditions on five transcripts at five different cell stages are shown compared to the L4440 empty vector RNAi control. At least 4 embryos were assayed for each data point. Standard deviations are shown as shaded ribbon regions. # indicates data analyzed in (E). **(E)** Distributions of *nos-2* mRNA cluster size under MEX-3, SPN-4 (ts), and dual MEX-3/SPN-4 depletion conditions at the 16 to 24-cell stage demonstrate decreased cluster sizes when compared to mock (L4440) depletion. Significance indicates p-values derived from multiple test corrected t-tests (0.005 > ** > 0.0005; 0.0005 > *** > 0.00005)

Curiously, we noticed transcripts did not mix homogenously within P granules but occupied discrete regions within granules depending on their type. For example, *clu-1* mRNA typically surrounded a *chs-1* mRNA core (Fig. 6). These observations are echoed by other reports of homo-typic mRNA spatial separation within germ granules [Eagle et al., 2018, Trcek et al., 2015] and suggest a complex organization to granules and the mRNAs they contain.

**Figure 6.**
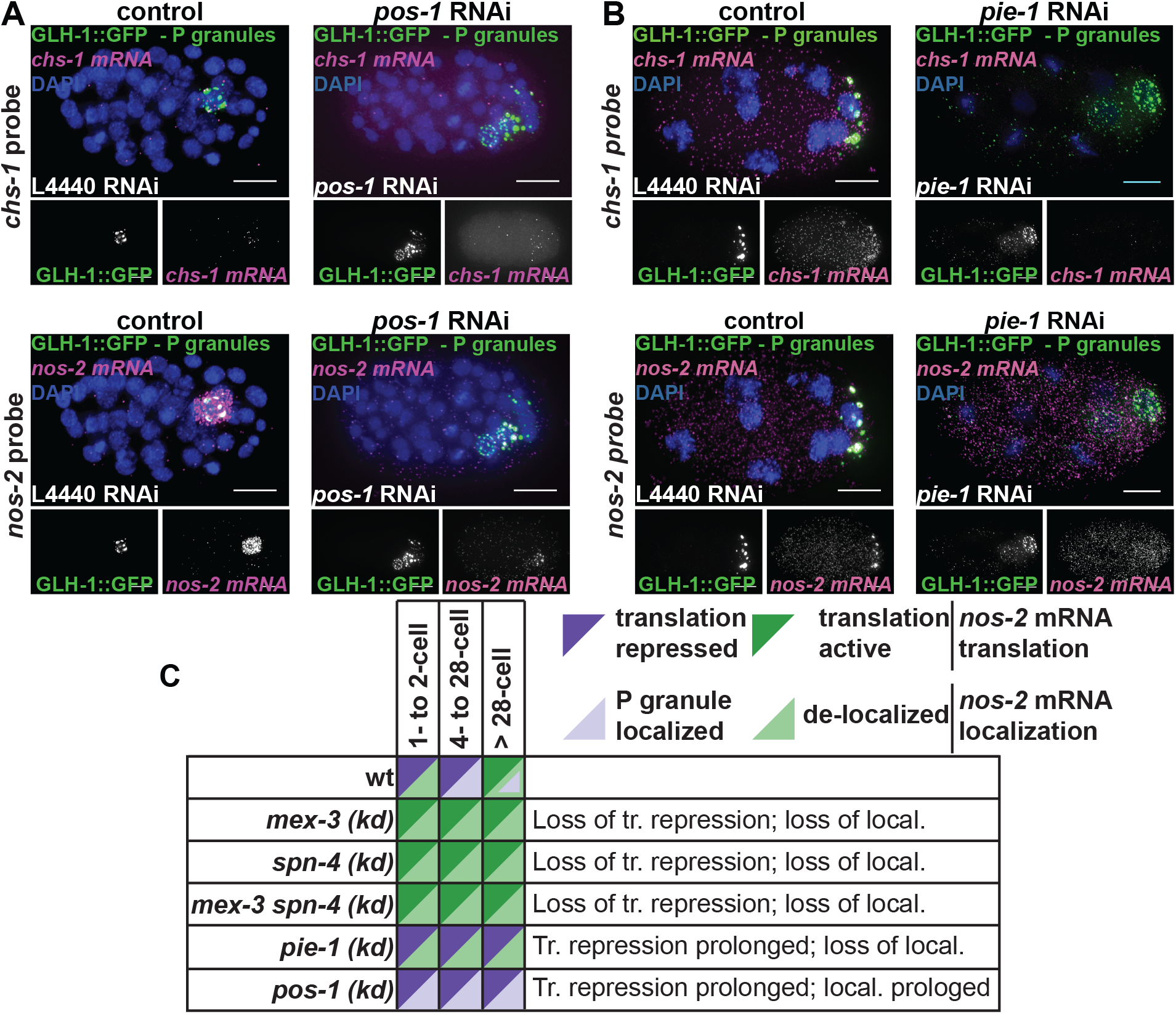
RBPs that regulate translation of NOS-2 differentially impact *nos-2* **mRNA subcellular localization**. **(A, B)** The impact of depleting POS-1 (A) or PIE-1 (B), two RBPs important for translation activation of *nos-2* mRNA at the 28-cell stage, was assayed. *chs-1* mRNA (magenta) and *nos-2* mRNA (magenta) were imaged in knock-down and control conditions using smFISH in a GLH-1∷GFP expressing strain. DNA was also stained with DAPI to illustrate developmental stage. The 28-cell stage is shown for *pos-1* RNAi conditions to illustrate the point in development when *nos-2* becomes translationally active. 8-cell stage embryos are shown for *pie-1* RNAi conditions to illustrate a stage when *nos-2* is still repressed. **(C)** Schematic showing a summary of phenotypes exhibited in knocking down RBPs that promote or inhibit *nos-2* translation and their impact on NOS-2 protein production (inferred from references) and *nos-2* mRNA localization (Fig. 5, Fig. 6).

### 3’UTRs were sufficient to direct mRNAs to P granules but not membranes

3’UTRs of transcripts have been implicated in driving subcellular localization of mRNAs in many organisms [Martin and Ephrussi, 2009]. To determine whether 3’UTRs of transcripts in our study were sufficient to direct mRNA localization, we appended 3’UTRs of interest onto *mNeonGreen* reporters expressed from the *mex-5* promoter in transgenic strains. We imaged *mNeonGreen* mRNA localization using *mNeonGreen* smFISH probes alongside probe sets for endogenous mRNA in the same embryos.

3’UTRs of *erm-1* and *imb-2* were not sufficient to drive mRNA subcellular localization. Endogenous *erm-1* and *imb-2* mRNAs localize to the cell or nuclear peripheries, respectively, but *mNeonGreen* mRNA appended with *erm-1* or *imb-2* 3’UTRs failed to recapitulate those patterns (Fig. 4A - 4D). However, the *imb-2* 3’UTR did show evidence of mRNA destabilization as *Pmex-5∷mNeonGreen∷imb-2 3’UTR* yielded fewer *mNeonGreen* mRNA than endogenous *imb-2* transcripts or *Pmex-5∷mNeonGreen∷erm-1 3’UTR* expressed under the same promoter. This suggests that sequences within the body of the *imb-2* mRNA and/or its successful localization are important for mRNA stability. Ultimately, we did not identify sequences within *erm-1* or *imb-2* mRNAs sufficient to direct transcript localization. Either the 5’ regions of the mRNA, the coding sequence of the mRNA, the full mRNA, a short N-terminal signal peptide, or some larger aspect of the translated protein direct mRNA localization.

In contrast, 3’UTRs of *chs-1*, *clu-1*, *cpg-2*, and *nos-2* were sufficient to direct *mNeonGreen* mRNA to P granules. Each of these *Pmex-5∷mNeonGreen∷3’UTR-of-interest* strains yielded *mNeonGreen* mRNA transcripts localized to P granules coincident with the localization of their endogenous mRNA (Fig. 4E - 4H, Fig. 7). The *chs-1* 3’UTR did exhibit some hallmarks of transcript destabilization given the comparative low abundance of *mNeonGreen∷chs-1 3’UTR* transcripts (Fig. 7A, B).

**Figure 7.**
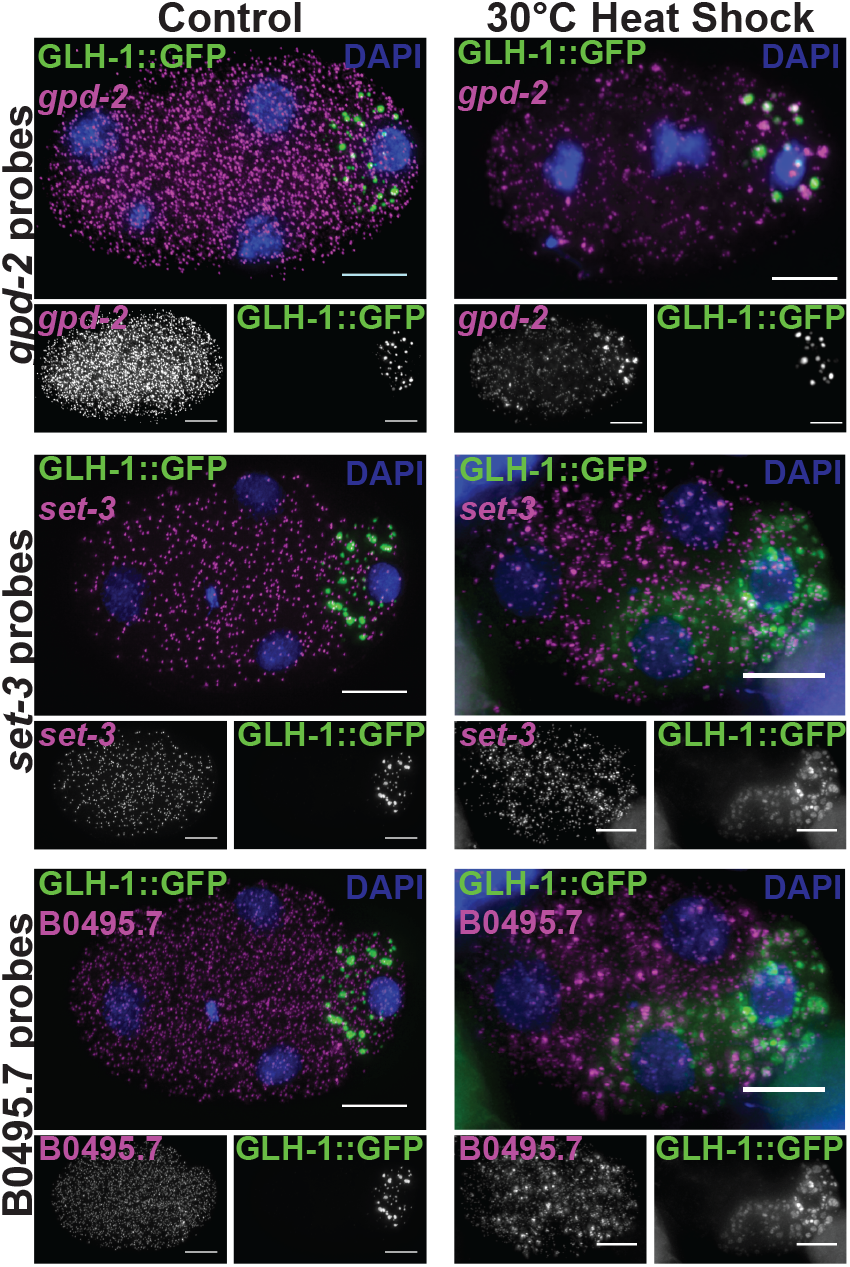
Homogenously distributed transcripts form clusters when subjected to heat shock stress. The transcripts *gpd-2, set-3*, and B0495.7 (magenta) are homogenously distributed in 4-cell embryos raised at 20°C (top). These transcripts become recruited to GLH-1∷GFP labeled P granules (green) and other uncharacterized mRNA clusters following a 25 minute 30°C heat shock (bottom). DNA was also stained with DAPI to illustrate developmental stage.

### RNA localization trends with translational status

NOS-2 protein is translationally repressed in germline and early embryonic stages before becoming translationally active in the P_4_ cell at the 28-cell stage with both repression and de-repression being mediated by *nos-2*’s 3’UTR [D’Agostino et al., 2006]. NEONGREEN protein under control of the *nos-2* 3’UTR in our study phenocopied this reported pattern (Fig. 8A). *neongreen* fused to 3’UTRs of other transcripts (*erm-1*, *imb-2*, *chs-1*, *clu-1*, or *cpg-2*) produced low levels of diffuse fluorescence, complicating interpretation of their translational status (Fig. 8B). GFP fusions to ERM-1, CHS-1, and CPG-2 proteins were more informative in illustrating the endogenous expression patterns of these proteins.

ERM-1∷GFP localized to the cell cortex throughout embryogenesis consistent with the role of the ERM-1 protein in linking the cortical actin cytoskeleton to the plasma membrane [Göbel et al., 2004, Van Fürden et al., 2004] (Fig. 9A). CHS-1 and CPG-2 play a more transient role in development, evidenced by GFP fusion reporters showing highest signal in the early cell stages (Fig. 9B, C). CHS-1 and CPG-2 work together to form two different layers of the trilaminar eggshell. CHS-1 encodes a multipass membrane protein that is activated and exocytosed upon fertilization to polymerize chitin forming the middle of the eggshell – the chitin layer – thereby blocking polyspermy [Maruyama et al., 2007, Olson et al., 2012]. CHS-1 enzymes then internalize stimulating exocytosis of CPG-1 and CPG-2 proteins that nucleate 5 and 34 chondroitin molecules respectively to form the inner eggshell layer – the CPG layer. mRNAs encoding both *chs-1* and *cpg-2* then decline as evidenced by our smFISH data. Indeed, a GFP fusion of CHS-1 show fluorescence at the 1-cell stage, but rapidly disappears thereafter (Fig. 9B). CPG-2 appears external to the cells and persists within the extracellular space but not within cells (Fig. 9C). Overall, transcripts undergoing active expression localized to the cytoplasm or membranes whereas mRNA transcripts whose expression was repressed, declining, or low tended to accumulate in P granules.

### Translational repressors of NOS-2 are required for mRNA degradation of multiple transcripts and are required for P granule localization of *nos-2* mRNA

*nos-2* is one of three *nanos*-related genes in the *C. elegans* genome and a member of the evolutionarily conserved *nanos* family. Similar to Drosophila *nanos* mRNA, *C. elegans nos-2* mRNA is contributed maternally, concentrates in the progenitor germ lineage, is translationally repressed in oocytes and during early embryogenesis, is translated with spatial specificity, and produces a protein whose final expression is restricted only to germ cells [Subramaniam and Seydoux, 1999]. *C. elegans nos-2* is required for proper development of the germ cells and is necessary with zygotically-expressed *nos-1* for germ cell proliferation. Translational repression of *nos-2* is coordinated by four sequential RNA Binding Proteins (RBPs) – OMA-1, OMA-2, MEX-3, and SPN-4 – that directly interact with the *nos-2* 3’UTR [D’Agostino et al., 2006, Jadhav et al., 2008] (Fig. 5A). In oocytes, OMA-1 and OMA-2 are redundantly required to repress translation through direct interactions with *nos-2*’s 3’UTR before they are degraded in the zygote. The RBPs MEX-3 and SPN-4 next take over. MEX-3 and SPN-4 repress *nos-2* translation throughout the embryo, with SPN-4 being most effective in posterior cells. MEX-3 and SPN-4 both interact with either of two directly repeated RNA sequences in the *nos-2* 3’UTR and function non-redundantly in the early embryo as RNAi or mutants of either results in premature translation of a *nos-2* reporter. This baton-passing of translational control has been documented for other maternally inherited transcripts including *zif-1* (an E3 ubiquitin ligase specific to somatic cells) [Oldenbroek et al., 2012] and *mom-2* (the Wnt ligand in P_2_) [Oldenbroek et al., 2013].

Though the requirement for OMA-1, OMA-2, MEX-3, and SPN-4 to repress translation of *nos-2* mRNA is clear, due to limitations of previous techniques, it is not known whether they are required to localize *nos-2* mRNA to P granules. To rectify this and to expand the question, we tested how depletion of these RBPs, individually or in combinations, impacted the abundance and/or localization of four clustered mRNA transcripts (*chs-1, clu-1, cpg-2*, and *nos-2*) (Fig. 5A). True to published reports, individual knockdowns of OMA-1 and OMA-2 had minimal phenotypes, but in combination yielded too few embryos to credibly test as development arrests during oogenesis [Detwiler et al., 2001, Shimada et al., 2002]. Depletion of MEX-3 and/or SPN-4 led to an overabundance of embryo-wide *chs-1, cpg-2*, and *nos-2* transcripts compared to mock RNAi control, suggesting that MEX-3 and SPN-4 have a direct or indirect role in mRNA degradation (Fig. 5B, C and Fig. 10). MEX-3 and SPN-4 are not required independently to accumulate *chs-1, clu-1*, or *cpg-2* mRNAs in P granules; however, double knockdown of MEX-3 and SPN-4 resulted in a loss of *chs-1* localization to P granules (Fig. 10). Only the localization of *nos-2* mRNA to P granules was severely disrupted by MEX-3 or SPN-4 loss independently as evidenced by the missing *nos-2* clusters in smFISH images (Fig. 5D, E) and corresponding decrease in the average number of mRNA molecules per cluster (Fig. 5C). Together, these findings suggest that MEX-3 and SPN-4 are required for both *nos-2*’s translational repression and P granule localization [D’Agostino et al., 2006, Jadhav et al., 2008]. Further, the role of MEX-3 and SPN-4 in RNA degradation is separable from their roles in mRNA localization to P granules as *chs-1, cpg-2*, and *nos-2* require MEX-3 and SPN-4 for RNA clearance whereas only *nos-2* and *chs-1* rely on them for P granule localization.

### RBPs that relieve NOS-2 translational repression impact *nos-2* localization differently

*nos-2* mRNA is translationally repressed in the germline, through fertilization, and is only released from repression at the 28-cell stage of development when NOS-2 protein is exclusively produced in the P_4_ cell [D’Agostino et al., 2006, Subramaniam and Seydoux, 1999, Tenenhaus et al., 2001]. *nos-2* mRNA localizes to P granules in the adult germline [Schisa et al., 2001], but appears distinct from P granules at the 1-cell and 2-cell stages (this study). Between the 4-cell and 28-cell stages, *nos-2* progressively re-accumulates into P granules reaching a maximum average density of 20 – 30 mRNA molecules per P granule prior to the 28-cell stage (Fig. 2, Fig. 2C). At the 28-cell stage of development when NOS-2 translation begins [Subramaniam and Seydoux, 1999], we observed *nos-2* mRNA becoming dispersed in the cytoplasm external to P granules (Fig. 6A). This could suggest that *nos-2* mRNA emerges from P granules when it becomes actively translated, supported by the fact that P granules are devoid of key ribosomal components required for translation [Schisa et al., 2001].

Given that loss of *nos-2* translational repression led to loss of *nos-2* mRNA localization within P granules (above, Fig. 5), we sought to determine the effects of prolonged *nos-2* translational repression. We imaged *nos-2* mRNA by smFISH under *pie-1* and *pos-1* RNAi knock-down conditions. PIE-1 and POS-1 RBPs both encode proteins that assist in relieving the translational repression of *nos-2* at the 28-cell stage [D’Agostino et al., 2006, Tenenhaus et al., 2001]. Interestingly, the two knock-down conditions yielded different results. Under *pos-1* RNAi, *nos-2* mRNA failed to appear in the cytoplasm and instead remained associated predominantly with P granules (Fig. 6A), correlating with its translationally inactive status. In contrast, depletion of PIE-1 had the opposite effect. PIE-1 is an RBP that plays a three-fold role contributing to *nos-2* stabilization, NOS-2 translational activation, and germline transcriptional repression [D’Agostino et al., 2006, Tenenhaus et al., 2001]. Upon disruption of PIE-1, *nos-2* mRNA molecules undergo progressive degradation in the P lineage due to the inappropriate transcription of somatic genes within the P lineage. If this degradation phenotype is abrogated by concurrent loss of *ama-1* (encoding RNA Polymerase II), *nos-2* mRNA molecules remain but in the absence of PIE-1 fails to produce NOS-2 protein, illustrating that PIE-1 also contributes to optimal translation of NOS-2 in the P lineage. Upon *pie-1* depletion, we confirmed premature *nos-2* mRNA degradation; however, we were surprised to see a complete loss of *nos-2* localization to P granules despite *nos-2* being translationally inactive at these stages [Tenenhaus et al., 2001]. (Fig. 6B). Initially, we suspected that P lineage identity was dysfunctional in these embryos leading to the loss of wild-type P granule function. However, P granules are clearly present in these embryos by GLH-1∷GFP marker proteins and they accumulate other mRNAs such as *clu-1* (Fig. 6, Fig. 11). Because *nos-2* mRNA is not translated upon *pie-1* disruption [Tenenhaus et al., 2001], this suggests that the translational repression of *nos-2* and its localization to P granules can be uncoupled, perhaps mimicking a somatic-cell-like state in the P lineage.

Taken together, RBP knockdown conditions that disrupt *nos-2* mRNA translational repression also disrupt *nos-2* mRNA P granule association (*mex-3* (RNAi) and *spn-4* (ts)). In contrast, an RBP knockdown condition that prolongs *nos-2* translational repression fails to release *nos-2* transcripts from P granules (*pos-1* (RNAi)). Therefore, the localization of *nos-2* mRNA in P granules is largely coincident with a translationally repressed state (Fig. 6C). It is not a perfect association, however. We observed several cases where *nos-2* mRNA remains translationally repressed even though they do not localize to P granules: 1) in 1-to 2-cell stage embryos where an abundance of cluster-independent *nos-2* mRNA are present throughout the embryo; 2) in somatic cells of the early embryo that contain numerous individual *nos-2* mRNA (up to ~80%); and 3) in *pie-1* mutants where *nos-2* fails to localize to P granules. These findings illustrate *nos-2* translational repression can occur independently of transcript localization and translational repression is not dependent on P granule residency. Further, it illustrates an order of operations in which translational repression precedes P granule localization during development.

### Disrupting translation promotes P granule localization

We speculated whether P granule localization was a natural consequence that befalls transcripts experiencing low rates of translation or complete repression. We were led to this hypothesis by several lines of evidence. *nos-2, chs-1*, and *cpg-2* mRNAs are either known or likely to be undergoing minimal translation in early embryogenesis and all localized to P granules in our studies. Further, P granules are depleted for rRNA, suggesting that protein synthesis does not occur within them [Schisa et al., 2001]. Finally, under heat, osmolarity, or oxidative stress, cells slow or stop translation and concurrently form stress granules (phase condensate compartments) that store up to 95% of untranslated transcripts [Khong et al., 2017], an example of low translational status resulting in mRNA sequestration into condensates.

To determine whether altering the translational status of mRNAs could change their localization within the cell, we disrupted translational initiation through heat exposure. Embryos exposed to 30°C for 25 minutes repress protein synthesis at the level of translational initiation. We observed that transcripts that are normally homogenously distributed throughout the cytoplasm and across the embryo coalesced into P granules stimulated by this heat stress. We observed this for three transcripts, *set-3 (SET domain containing), gpd-2 (Glycerol-3-Phosphate Dehydrogenase)*, and B0495.7 *(predicted metalloprotease)* (Fig. 7). Therefore, loss of protein synthesis was able to ectopically stimulate otherwise homogenous mRNA transcripts to accumulate in P granules. This further indicates that P granule accumulation is a downstream step of translational repression and that P granule accumulation requires translational repression or low levels of translation to stimulate the localization of a transcript.

## Discussion

### Translational repression of mRNA is necessary and sufficient for P granule localization

In this study, we report several maternally inherited mRNAs with subcellular localization in early *C. elegans* embryos. In many cases, these patterns of localization are linked to the RNA’s translational status, particularly in the case of P granule-associated transcripts that have low rates of translation or are fully repressed. We envisioned translationally repressed transcripts could accumulate in P granules by different mechanisms. Either mRNAs are actively brought to P granules for the purpose of translational repression (due to the paucity of ribosomal components there), or they are translationally repressed in the cytoplasm leading to accumulation in P granules as a downstream step. In the case of *nos-2* our evidence supports the second model in which translational repression precedes and directs P granule localization. Though *nos-2* mRNA translational repression and P granule localization strongly correlated, we observed situations where these properties were uncoupled (1-cell stage, somatic cells, and upon *pie-1* depletion). In these cases, *nos-2* translational repression did not depend on localization to P granules. Further, translational down-regulation occurred before P granule localization. We showed that transcripts can ectopically localize into P granules upon disruption of translational initiation, illustrating that translational repression is sufficient to direct P granule localization. Together, these findings support the model that mRNAs of low translational status accumulate in P granules as a downstream step.

### P granules functionally echo stress granules and P-bodies by accumulating transcripts of low translational status

mRNAs that localize to P granules can still be observed as individuals within the cytoplasm. Indeed, from 7% (*clu-1*, 26-48-cell stage) to 53% (*clu-1*, 8-cell stage) of total mRNAs of different species localized to clusters as opposed to being present as individuals. This echoes situations where stress-induced translational disruption promotes transcripts to move into stress granules. In those cases, 10% of bulk mRNA and up to 95% of specific transcripts move into stress granules only returning to the cytoplasm after the stress has passed [Khong et al., 2017]. Though stress granules and germ granules (like P granules) are distinct, they appear to have some functionality in common.

Future studies will uncover more detailed causal relationships between translational regulatory control and the subcellular localization patterns we have discovered. Indeed, the early embryo presents us with a unique opportunity to uncover novel mechanisms of post-transcriptional gene regulatory control as our observations of existing mRNA populations are not confounded by *de novo* transcription.

### Different transcripts accumulate in P granules through different mechanisms

We identified six new genes from the list of P_1_-enriched transcripts [Osborne Nishimura et al., 2015] that yielded clustered patterns of mRNA localization. We select three of these (*chs-1, clu-1*, and *cpg-2*) for further study. All three transcripts localized to P granules in 3’UTR-dependent manners. However, these transcripts did not rely on the same RBPs (MEX-3, SPN-4, and PIE-1) to the same extent as did *nos-2* for localization into granules. What, then, directs *clu-1* and *cpg-2* to P granules? What drives the differences between *chs-1* and *nos-2*’s dependence on MEX-3 and SPN-4? The answer may lie in their biology. CHS-1 and CPG-2 are translationally activated by fertilization but they decrease in both mRNA abundance and in protein production shortly thereafter. Therefore, whether translation is repressed temporarily as in the case of *nos-2* or permanently and followed by degradation as in the case of *chs-1* or *cpg-2* it is possible that minimal translational activity can generally lead to P granule accumulation. Because each transcript has varied dependence on MEX-3, SPN-4, and PIE-1 for their localization to P granules, it suggests different sets of RBPs interpret the 3’UTR-directed sequence information encoding their fates in different manners.

### mRNA degradation plays a role in shaping transcript localization patterns

Transcripts of *chs-1, clu-1, cpg-2*, and *nos-2* accumulate in the P granules of progenitor germ cells at the same time they disappear from somatic cells. These linked mechanisms concentrate transcripts down the P lineage. All transcripts tested required MEX-3 and SPN-4 for degradation in somatic cells, yet *nos-2* specifically required both of these RBPs for strong accumulation in P granules. *chs-1* and *cpg-2* mRNAs were also altered in their localization upon loss of these RBPs, but not to the same extent as *nos-2*. Together these findings suggest a mechanism in which P granule and/or P lineage localization of these transcripts protects mRNAs from MEX-3 and SPN-4-dependent degradation pathways while promoting their recruitment to P granules. This mechanism of local protection coupled to generalized degradation has also been evoked to explain how Drosophila *nanos* concentrates at posterior regions of the embryonic syncytium in that specie [Lasko, 2012]. Similarly, we found the 3’UTR of *imb-2* fused to *mNeonGreen* elicited *mNeonGreen* mRNA decay suggesting that *imb-2* localizes to nuclei by a 3’UTR-independent mechanism that protects it from its own 3’UTR-dependent degradation. Together, these findings illustrate how RNA degradation can carve out cell-specific patterning and how subcellular localization can protect RNAs to preserve them in specific regions of the cell and embryo.

Of the 8 P_1_-enriched transcripts we imaged using smFISH, all overlapped with P granule markers. We are interested to determine the translational states of more of these transcripts to determine whether the correlation between P granule localization and low translational status holds. We are intrigued by the possibility that translational status directs P granule residency, and P granule residency, in turn, directs enrichment down the P lineage. This explains how mRNAs may be retained and concentrated in specific lineages even in the absence of *de novo* transcription.

### Peripheral transcripts often encode membrane-associated proteins

Half of the anterior, AB-enriched transcripts we surveyed by smFISH accumulated at the cell periphery. These transcripts included *erm-1 (Ezrin/Radixin/Moesin), lem-3 (LEM domain protein), ape-1 (APoptosis Enhancer)*, and *tes-1 (TEStin homolog)*. ERM-1 proteins also localize to the apical plasma membrane where they link the cortical actin cytoskeleton to the plasma membrane, suggesting a functional linkage between mRNA and protein localization [Van Fürden et al., 2004]. Further, LEM-3 localizes to apical membranes, the midbody, and cytoplasmic foci [Dittrich et al., 2012]. The localizations of APE-1 and TES-1 are currently uncharacterized, but these proteins harbor domains associated with membrane localization [Bennett and Baines, 2001, Sweede et al., 2008]. In addition, we discovered that the symmetrically distributed (at the 2-cell stage) transcript *imb-2 (IMportin Beta)* localized preferentially at nuclear membranes, the same localization where the protein it encodes functions [Putker et al., 2013]. The concordance between localization of mRNA and the proteins they encode suggest that either the transcripts are directed to membranes for the purpose of local translation or they are passively dragged along behind the growing peptide as it localizes to its final destination. Current genomics assays have illustrated that mRNAs can associate with membranes and/or the ER in both translationally-dependent and -independent ways, suggesting both models are possible.

### Computational toolkit for assessing mRNA patterning

We developed techniques to quantify and computationally describe the subcellular patterns we observed. By identifying individual mRNAs in relationship to cellular landmarks or in relation to one another, we could quantitate our findings and illustrate differences. Doing so, we were able to estimate the number of mRNAs of different types within P granules. We found that transcripts in P lineage cells were often present at 8 – 12 transcripts per granule but *nos-2* mRNA in particular could accumulate to higher titers (>20 molecules per granule) just prior to the onset of *nos-2* translation, indicating a high concentration of *nos-2* mRNA in P granules at that stage. Overall, we expect our approach of determining the proximity of RNAs in relation to cellular landmarks and deconvolving overlapping smFISH signals will be of broad interest.

### mRNA localization is a widespread feature of cell biology

Diverse examples of transcript-specific mRNA localization have been described across the tree of life ranging from bacteria [Fei and Sharma, 2018] to humans [Khalil et al., 2018]. Major inroads in understanding mRNA localization patterns and their mechanisms and functions have been contributed by a variety of organisms including yeast (*ASH-1* mRNA defines mother versus bud cell identity), Drosophila oocytes and early embryos (*oskar, bicoid, gurken*, and *nanos* impact cell fate and embryonic development), and mammalian neurons (*β-actin* assists in axonal guidance). In almost all of these systems, localization of mRNAs correlating with modes of post-transcriptional regulatory control. Initially, the oocyte and neurons were hypothesized to represent special cases where mRNA localization played an augmented role due to the multi-nucleate nature of the Drosophila syncytium or the complex morphology of a nerve cell. However, recent advances in mRNA imaging and proximity labeling are starting to suggest mRNA localization and its control is more widespread. A new perspective is emerging to encompass mRNA localization control as a general feature of cell biology.

## Materials and Methods

### *C. elegans* maintenance

*C. elegans* strains were maintained using standard procedures [Brenner, 1974]. Worms were grown at 20°C and reared on Nematode Growth Medium (NGM: 3 g/L NaCl; 17 g/L agar; 2.5 g/L peptone; 5 mg/L cholesterol; 1 mM CaCl2; 1 mM MgSO4; 2.7 g/L KH2PO4; 0.89 g/L K2HPO4). *C. elegans* strains generated in this study were derived from the standard laboratory strain, Bristol N2. Strains used in this study are listed in Supplementary Table 1.

### 3’UTR Reporter Constructs

The plasmid pMTNCSU7 was generated to express mNeonGreen as an N-terminal fluorescent reporter. Starting with a *Pmex-5∷neongreen∷neg-1∷neg-1-3’UTR* plasmid derived from the MosSCI-based plasmid pCFJ150, we replaced the *neg-1* sequences with an NheI/BglII/EcoRV multiple cloning site using inverse PCR. 3’UTRs were PCR amplified and cloned into the NheI site of pMTNCSU7 using Gibson cloning (NEB) to create pDMP45 *(Pmex-5∷mNeonGreen∷nos-2 3’UTR)*, pDMP47 *(Pmex-5∷mNeonGreen∷cpg-2 3’UTR)*, pDMP48 *(Pmex-5∷mNeonGreen∷chs-1 3’UTR)*, pDMP91 *(Pmex-5∷mNeonGreen∷clu-1 3’UTR)*, pDMP111 *(Pmex-5∷mNeonGreen∷imb-2 3’UTR)*, and pDMP112 *(Pmex-5∷mNeonGreen∷erm-1 3’UTR)*. Plasmids used in this study are listed in Supplementary Table 2. Primers used for 3’UTR amplification can be found in Supplementary Table 3.

### *C. elegans* Single-Copy Transgenesis by CRISPR

*Pmex-5∷mNeonGreen∷3UTR* strains were generated from N2 worms by CRISPR targeting to the ttTi5605 MosSCI site [Dickinson et al., 2013]. Guide RNA targeting the ttTi5605 MosSCI site and Cas9 protein were co-expressed from the plasmid pDD122 while plasmids pDMP45, pDMP47, pDMP48, pDMP91, pDMP111, and pDMP112 were used as repair templates. Three vectors containing mCherry tagged pGH8 (*Prab-8∷mCherry* neuronal co-injection marker), pCFJ104 (*Pmyo-3∷mCherry* body wall muscle co-injection marker), and pCFJ90 (*Pmyo-2∷mCherry* pharyngeal co-injection marker) as well as one containing the heat-shock activated PEEL-1 counter-selectable marker (pMA122) were coinjected. mNeonGreen and mCherry positive animals were identified as F1 progeny and singled to new plates until starvation. Starved plates were then subjected to a 4-hour incubation at 34°C to counter-select, followed by an overnight recovery at 25°C. Plates were then screened for living worms that did not express the mCherry coinjection markers. Worms that showed no fluorescence from the presence of extrachromosomal arrays were singled to establish lines, which were confirmed for single-copy insertion by PCR using the primers in Supplementary Table 3.

### smFISH

single molecule Fluorescence *In Situ* Hybridization (smFISH) was performed based on the TurboFish protocol with updates specific to *C. elegans* and using new Biosearch reagents [Femino et al., 1998, Osborne Nishimura et al., 2015, Raj et al., 2008, Raj and Tyagi, 2010, Shaffer et al., 2013]. Custom Stellaris FISH Probes were designed against target transcripts (Supplementary Table 4) by utilizing the Stellaris RNA FISH Probe Designer (Biosearch Technologies, Inc., Petaluma, CA) available online at www.biosearchtech.com/stellarisdesigner (version 4.2). The embryos were hybridized with Stellaris RNA FISH Probe sets labeled with Cal Fluor 610 or Quasar 670 (Biosearch Technologies, Inc.), following the manufacturer’s instructions available online at www.biosearchtech.com/stellarisprotocols. Briefly, young adult worms were bleached for embryos, suspended in 1 ml −20°C methanol, quickly vortexed, and freeze cracked in liquid nitrogen. Embryos were stored in methanol at −20°C for 1-24 hours. After fixation, embryos were equilibrated briefly in Stellaris Wash Buffer A (Biosearch, SMF-WA1-60) before hybridization in 100 *μ*l Stellaris Hybridization buffer (Biosearch, SMF-HB1-10) containing 10% formamide and 50 picomoles of each primer set. The hybridization reaction was incubated at 37°C overnight. Hybridized embryos were then washed twice for 30 min in Stellaris Wash Buffer A with the second wash containing 1 *μ*g/ml of DAPI. Following counterstaining, a final wash in Stellaris Wash Buffer B (Biosearch, SMF-WB1-20) was carried out before storage with n-propyl gallate antifade (10 ml 100% glycerol, 100 mg N-propyl gallate, 400 *μ*l 1M Tris pH 8.0, 9.6 ml DEPC treated H2O) prior to slide preparation. Embryos were mounted based on original descriptions in [Ji and van Oudenaarden, 2012], using equal volumes hybridized embryos resuspended in N-propyl gallate antifade and Vectashield antifade (Vector Laboratories, H-1000). smFISH image stacks were acquired on a Photometrics Cool Snap HQ2 camera using a DeltaVision Elite inverted microscope (GE Healthcare), with an Olympus PLAN APO 60X (1.42 NA, PLAPON60XOSC2) objective, an Insight SSI 7-Color Solid State Light Engine, and SoftWorx software (Applied Precision) using 0.2 *μ*m z-stacks. Representative images were deconvolved using Deltavision (SoftWorx) deconvolution software. Images were further processed using FIJI [Schindelin et al., 2012]. Initial characterization of subcellular localization for the transcripts *erm-1, imb-2, chs-1, clu-1, cpg-2*, and *nos-2* was performed in conjunction with the homogenous transcript *set-3* as a negative control for subcellular localization (Data not shown).

### smiFISH

single-molecule inexpensive FISH was performed as in [Tsanov et al., 2016] using FLAPY primary probe extensions and secondary probes. Briefly, between 12 and 24 primary probes were designed using Oligostan [Tsanov et al., 2016] and ordered in 25 nmol 96-well format from IDT diluted to 100 *μ*M in IDTE buffer pH 8.0. Secondary FLAPY probes were ordered from Stellaris LGC with dual 5’ and 3’ fluorophore labeling using either Cal Fluor 610 or Quasar 670. Individual probes were combined to a final concentration of 0.833 *μ*M. 2 *μ*l of primary probe mixture were mixed with 1 *μ*l 50 *μ*M FLAPY secondary probe, 1 *μ*l NEB buffer 3, and 6 *μ*l DEPC treated H2O. The primary and secondary probe mixtures were then incubated in a thermocycler at 85°C for 3 min., 65°C for 3 min., and 25°C for 5 min. to anneal. 2 *μ*l of annealed probe mixtures were then used as normal smFISH probe sets as above. smiFISH probe sequences are listed in Supplementary Table 4.

### smFISH plus Immunofluorescence

smFISH combined with immunofluorescence was performed similarly to smFISH with slight modifications. N2 and DUP98 *patr-1(sam50[patr-1∷GFP∷3xLAG])II* [Andralojc et al., 2017] embryos were harvested as above with the exception that they were resuspended in methanol, freeze cracked in liquid nitrogen for 1 min., and transferred to acetone after ~5 min. total in methanol. Embryos were then incubated in acetone for 25 min. before proceeding to hybridization/immunofluorescence. smFISH was then performed as above with the exception that 2.37 *μ*g/ml Janelia Fluor 549 (Tocris, Cat. No. 6147) conjugated anti-GFP nanobody (Chromotek, gt-250) was incubated with the embryos overnight in hybridization buffer.

### Initial quantification of smFISH micrographs

Initial characterization of mRNA counts from smFISH micrographs were performed using a standard FISH-quant [Mueller et al., 2013] analysis. Briefly, embryos were manually outlined, 3D LoG filtered using default FISH-quant parameters (Size = 5, Standard deviation = 1), and spots were pre-detected using a local maximum fitting, and RNAs were detected using a manually determined image-dependent intensity and quality threshold with sub-region fitting of 2 pixels in the x and y axes and 3 pixels in the z axis.

Post-processing to calculate the different location metrics was performed as described below with custom written Matlab and Python code. The Python code is implemented as plugins for the image processing platform ImJoy [Ouyang et al., 2019]. Source code and detailed description are provided here http://github.com/muellerflorian/parker-rna-loc-elegans

### Quantification of Cortical RNA Localization

Quantification of transcript localization to the cell cortex was performed using the web application ImJoy [Ouyang et al., 2019]. RNAs were first detected as above using FISH-quant. Individual cell outlines were then manually annotated in FIJI for each Z-stack in the micrograph, excluding the uppermost and lowermost stacks where cells are flattened against the slide or coverslip. The distance of each RNA was then measured from the nearest annotated membrane and binned in 10 *μ*m increments. Total number of RNAs per bin were then normalized by the volume of the concentric spheres they occupied. After this normalization, values larger than 1 indicate that for this distance more RNAs are found compared to a randomly distributed sample.

### Quantification of Nuclear Peripheral RNA Localization

Quantification of transcript localization to the nuclear periphery was also performed using the ImJoy. RNAs were first detected as above using FISH-quant. Embryos were then manually outlined to create an upper limit for RNA distance from the nucleus. Individual nuclei were then annotated by binarizing DAPI micrographs to create a nuclear mask. The distance of each RNA was then measured from the nearest annotated nuclear membrane and binned in 10 *μ*m increments. Negative distance indicates positioning within the nuclear mask. Total number of RNAs per bin were then normalized for volume as described above for cell membrane localization.

### Quantification of RNA Clustering

Detection of RNA molecules was performed in the 3D image stacks with FISH-quant [Mueller et al., 2013]. Positions of individual RNA molecules within dense clusters, were determined with a recently developed approach using the signal of isolated RNAs to decompose these clusters [Samacoits et al., 2018]. Post-processing to calculate the different location metrics was performed as described below with custom written Matlab and Python code. The Python code is implemented in user-friendly plugins for the image processing platform ImJoy [Ouyang et al., 2019]. Source code and all scripts used for analysis and figure generation are available here http://github.com/muellerflorian/parker-rna-loc-elegans

To quantify the number of individual mRNAs in mRNA clusters, the total number of clusters per embryo, and the fraction of mRNAs in clusters a custom MATLAB script was implemented. FISH-quant detection settings were used to identify candidate mRNA clusters from smFISH micrographs using a Gaussian Mixture Model (GMM). The GMM differentiates independent, single mRNAs from groups of clustered mRNAs by probabilistically fitting a predicted RNA of average intensity and size over each FISH-quant detected RNA. GMM fitting then provided coordinates of both independent RNAs and the modeled coordinates of each RNA that composes a cluster. The decomposed coordinates of each RNA in the embryo were then used by a Density-Based Spatial Clustering of Applications with Noise (DBSCAN) algorithm to quantitatively analyze cluster size and number.

### Quantifying RNA cluster overlap with GLH-1∷GFP

To determine the degree of overlap between RNA clusters and P granules labeled with GLH-1∷GFP a hybrid Matlab-ImJoy pipeline was implemented. RNA clusters were identified as described above. The occupied volume of these clusters in the image was calculated as the convex hull around all RNAs positions within a cluster with the SciPy function ConvexHull. Location of P granules was determined in 3D with a Laplacian of Gaussian (LoG) blob detection method (with the scikit-image function blog log). RNA clusters and P granules were considered to co-localize when their 3D volumes at least partly overlap. This allowed quantification of the number of independent P granules, RNA clusters, and RNA clusters that overlap with P granules.

### RNAi Feeding for smFISH Microscopy

dsRNA feeding was executed as previously described [Sawyer et al., 2011]. Mixed stage worms were bleached to harvest and synchronize embryos. Harvested embryos were deposited on RNAi feeding plates and grown at 25°C until gravid. Embryos were harvested and smFISH was conducted. For each gene targeted by RNAi, we performed at least three independent replicates of feeding and smFISH using L4440 empty vector as a negative control and *pop-1* RNAi as a 100% embryonic lethal positive control. For experiments using the *spn-4* temperature sensitive allele, *spn-4(or191) V*, worms were grown at 15°C until gravid, bleached for embryos, and split into 15°C negative control and 25°C query conditions while plating on L4440, *mex-3*, or *pop-1* RNAi conditions.

## Acknowledgments

We thank Michael Boxem, Tom Evans, Dan Dickinson, Chih Yung Lee, Chris Link, Brooke Montgomery, Tai Montgomery, Ari Pani, Sean Ryder, Geraldine Seydoux, Timothy Stasevich, Dustin Updike, Ning Zhao, and WormBase for protocols, equipment, reagents, strains, advice, productive discussions, feedback regarding the manuscript, and commentary. Some strains were provided by the CGC, which is funded by NIH Office of Research Infrastructure Programs (P40 OD010440).

## Funding

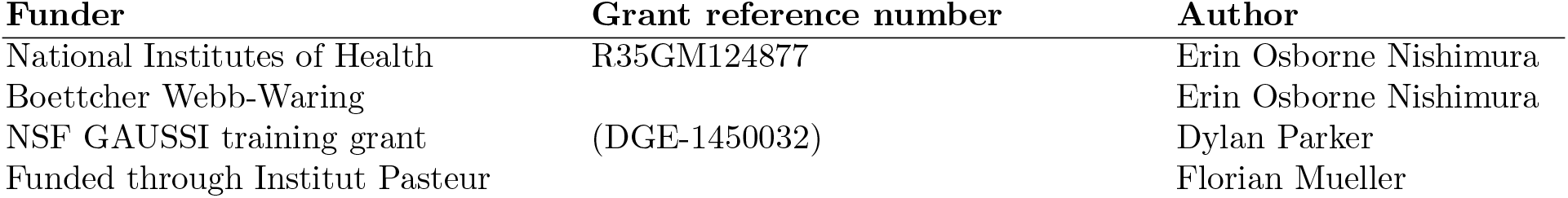

## Author Contributions

Dylan M. Parker, conceptualization, data collection, formal analysis, investigation, methodology, writing – original draft, writing – review and editing, funding acquisition; Lindsay P. Winkenbach, data collection, methodology, writing – review and editing; Samuel P. Boyson – data collection, methodology; Matthew Saxton – data collection; Camryn Daidone – data collection; Zainab Al Mayazdeh – data collection; Marc T. Nishimura – data collection, methodology, conceptualization; Florian Mueller – data curation, software, conceptualization, investigation; Erin Osborne Nishimura – conceptualization, data collection, methodology, formal analysis, funding acquisition, writing – original draft, writing – review and editing.

## References

Andralojc, K. M., Campbell, A. C., Kelly, A. L., Terrey, M., Tanner, P. C., Gans,I. M., Senter-Zapata, M. J., Khokhar, E. S., and Updike, D. L. (2017). ELLI-1, a novel germline protein, modulates RNAi activity and P-granule accumulation in *Caenorhabditis elegans*. PLoS Genetics, 13:(2)1–20.

Batchelder, C., Dunn, M. A., Choy, B., Suh, Y., Cassie, C., Shim, E. Y., Shin, T. H., Mello, C., Seydoux, G., and Blackwell, T. K. (1999). Transcriptional repression by the *Caenorhabditis elegans* germ-line protein PIE-1. Genes and Development, 13:(2)202–212.

Bennett, V. and Baines, A. J. (2001). Spectrin and ankyrin-based pathways: Metazoan inventions for integrating cells into tissues. Physiological Reviews, 81:(3)1353–1392.

Brangwynne, C. P., Eckmann, C. R., Courson, D. S., Rybarska, A., Hoege, C., Gharakhani, J., Jülicher, F., and Hyman, A. A. (2009). Germline P granules are liquid droplets that localize by controlled dissolution/condensation. Science, 5(JUNE):1729–1732.

Brenner, S. (1974). The genetics of *Caenorhabditis elegans*. Genetics, 77:(1)71–94.

Campbell, A. C. and Updike, D. L. (2015). CSR-1 and P granules suppress sperm-specific transcription in the *C. elegans* germline. Development (Cambridge), 142:(10)1745–1755.

D’Agostino, I., Merritt, C., Chen, P.-l., Seydoux, G., and Subramaniam, K. (2006). Translational repression restricts expression of the *C. elegans* Nanos homolog NOS-2 to the embryonic germline. Developmental Biology, 292:(1)244–252.

DeRenzo, C., Reese, K. J., and Seydoux, G. (2003). Exclusion of germ plasm proteins from somatic lineages by cullin-dependent degradation. Nature, 424:(6949)685–689.

Detwiler, M. R., Reuben, M., Li, X., Rogers, E., and Lin, R. (2001). Two zinc finger proteins, OMA-1 and OMA-2, are redundantly required for oocyte maturation in *C. elegans*. Developmental Cell, 1:(2)187–199.

Dickinson, D. J., Ward, J. D., Reiner, D. J., and Goldstein, B. (2013). Engineering the *Caenorhabditis elegans* genome using Cas9-triggered homologous recombination. Nature Methods, 10:(10)1028–1034.

Dittrich, C. M., Kratz, K., Sendoel, A., Gruenbaum, Y., Jiricny, J., and Hengartner, M. O. (2012). Lem-3 - a lem domain containing nuclease involved in the dna damage response in *C. elegans*. PLoS ONE, 7(2).

Eagle, W. V., Yeboah-Kordieh, D. K., Niepielko, M. G., and Gavis, E. R. (2018). Distinct cis-acting elements mediate targeting and clustering of *Drosophila* polar granule mRNAs. Development (Cambridge), 145(22).

Elewa, A., Shirayama, M., Kaymak, E., Harrison, P. F., Powell, D. R., Du, Z., Chute, C. D., Woolf, H., Yi, D., Ishidate, T., Srinivasan, J., Bao, Z., Beilharz, T. H., Ryder, S. P., and Mello, C. C. (2015). POS-1 promotes endo-mesoderm development by inhibiting the cytoplasmic polyadenylation of *neg-1* mRNA. Developmental Cell, 34:(1)108–118.

Ester, M., Kriegel, H. P., Sander, J., & Xu, X. (1996). A Density-Based Algorithm for Discovering Clusters in Large Spatial Databases with Noise. Proceedings of the second International Conference on Knowledge Discovery and Data Mining, pages 226–231.

Fei, J. and Sharma, C. M. (2018). RNA localization in Bacteria. Microbiology Spectrum, 6(5).

Femino, A. M., Fay, F. S., Fogarty, K., and Singer, R. H. (1998). Visualization of single RNA transcripts in situ. Science, 280:(5363)585–590.

Fields, S. D., Conrad, M. N., and Clarke, M. (1998). The *S. cerevisiae* CLU1 and *D. discoideum* cluA genes are functional homologues that influence mitochondrial morphology and distribution. Journal of Cell Science, 111:(12)1717–1727.

Gallo, C. M., Munro, E., Rasoloson, D., Merritt, C., and Seydoux, G. (2008). Processing bodies and germ granules are distinct RNA granules that interact in *C. elegans* embryos. Developmental Biology, 323:(1)76–87.

Gallo, C. M., Wang, J. T., Motegi, F., and Seydoux, G. (2010). Cytoplasmic partitioning of P granule components is not required to specify the germline in *C. elegans*. Science, 330:(6011)1685–1689.

Ghosh, D. and Seydoux, G. (2008). Inhibition of transcription by the *Caenorhabditis elegans* germline protein PIE-1: Genetic evidence for distinct mechanisms targeting initiation and elongation. Genetics, 178:(1)235–243.

Gibert, M., Starck, J., and Beguet, B. (1984). Role of the gonad cytoplasmic core during oogenesis of the nematode *Caenorhabditis elegans*. In Biology of the Cell, pages 77–85. Wiley.

Göbel, V., Barrett, P. L., Hall, D. H., and Fleming, J. T. (2004). Lumen morphogenesis in *C. elegans* requires the membrane-cytoskeleton linker *erm-1*. Developmental Cell, 6:(6)865–873.

Guven-Ozkan, T., Robertson, S. M., Nishi, Y., and Lin, R. (2010). *zif-1* translational repression defines a second, mutually exclusive OMA function in germline transcriptional repression. Development, 137:(20)3373–3382.

Hamm, D. C. and Harrison, M. M. (2018). Regulatory principles governing the maternal-to-zygotic transition: Insights from *Drosophila melanogaster*. Open Biology, 8(12).

Hashimshony, T., Feder, M., Levin, M., Hall, B. K., and Yanai, I. (2015). Spatiotemporal transcriptomics reveals the evolutionary history of the endoderm germ layer. Nature, 519:(7542)219–222.

Hashimshony, T., Wagner, F., Sher, N., and Yanai, I. (2012). CEL-Seq: single-cell RNA-Seq by multiplexed linear amplification. Cell Reports, 2:(3)666–673.

Hird, S. N., Paulsen, J. E., and Strome, S. (1996). Segregation of germ granules in living *Caenorhabditis elegans* embryos: Cell-type-specific mechanisms for cytoplasmic localisation. Development, 122:(4)1303–1312.

Jadhav, S., Rana, M., and Subramaniam, K. (2008). Multiple maternal proteins coordinate to restrict the translation of *C. elegans nanos-2* to primordial germ cells. Development, 135:(10)1803–1812.

Ji, N. and van Oudenaarden, A. (2012). Single-molecule fluorescent in situ hybridization (smFISH) of *C. elegans* worms and embryos. WormBook: the online review of C. elegans biology, pages 1–16.

Jud, M., Razelun, J., Bickel, J., Czerwinski, M., and Schisa, J. A. (2007). Conservation of large foci formation in arrested oocytes of *Caenorhabditis* nematodes. Development Genes and Evolution, 217:(3)221–226.

Kawasaki, I., Amiri, A., Fan, Y., Meyer, N., Dunkelbarger, S., Motohashi, T., Karashima, T., Bossinger, O., and Strome, S. (2004). The PGL family proteins associate with germ granules and function redundantly in *Caenorhabditis elegans* germline development. Genetics, 167:(2)645–661.

Khalil, B., Morderer, D., Price, P. L., Liu, F., and Rossoll, W. (2018). mRNP assembly, axonal transport, and local translation in neurodegenerative diseases. Brain Research, 1693:75–91.

Khong, A., Matheny, T., Jain, S., Mitchell, S. F., Wheeler, J. R., and Parker, R. (2017). The stress granule transcriptome reveals principles of mRNA accumulation in stress granules. Molecular Cell, 68:(4)808–820.e5.

Lasko, P. (2012). mRNA localization and translational control in *Drosophila* oogenesis. Cold Spring Harbor Perspectives in Biology, 4:(10)1–15.

Marnik, E. A. and Updike, D. L. (2019). Membraneless organelles: P granules in *Caenorhabditis elegans*. Traffic, 20:(6)373–379.

Martin, K. C. and Ephrussi, A. (2009). mRNA Localization: Gene expression in the spatial dimension. Cell, 136:(4)719–730.

Maruyama, R., Velarde, N. V., Klancer, R., Gordon, S., Kadandale, P., Parry, J. M., Hang, J. S., Rubin, J., Stewart-Michaelis, A., Schweinsberg, P., Grant, B. D., Piano, F., Sugimoto, A., and Singson, A. (2007). EGG-3 regulates cell-surface and cortex rearrangements during egg activation in *Caenorhabditis elegans*. Current Biology, 17:(18)1555–1560.

Merritt, C., Rasoloson, D., Ko, D., and Seydoux, G. (2008). 3’UTRs are the primary regulators of gene expression in the *C. elegans* germline. Current Biology, 18:(19)1476–1482.

Mueller, F., Senecal, A., Tantale, K., Marie-Nelly, H., Ly, N., Collin, O., Basyuk, E., Bertrand, E., Darzacq, X., and Zimmer, C. (2013). FISH-quant: automatic counting of transcripts in 3D FISH images. Nature Methods, 10:(4)277–278.

Oldenbroek, M., Robertson, S. M., Guven-Ozkan, T., Gore, S., Nishi, Y., and Lin, R. (2012). Multiple RNA-binding proteins function combinatorially to control the soma-restricted expression pattern of the E3 ligase subunit ZIF-1. Developmental Biology, 363:(2)388–398.

Oldenbroek, M., Robertson, S. M., Guven-Ozkan, T., Spike, C., Greenstein, D., and Lin, R. (2013). Regulation of maternal Wnt mRNA translation in *C. elegans* embryos. Development (Cambridge), 140:(22)4614–4623.

Olson, S. K., Greenan, G., Desai, A., Müller-Reichert, T., and Oegema, K. (2012). Hierarchical assembly of the eggshell and permeability barrier in *C. elegans*. Journal of Cell Biology, 198:(4)731–748.

Osborne Nishimura, E., Zhang, J. C., Werts, A. D., Goldstein, B., and Lieb, J. D. (2015). Asymmetric transcript discovery by RNA-seq in *C. elegans* blastomeres identifies *neg-1*, a gene important for anterior morphogenesis. PLoS Genetics, 11:(4)1–29.

Ouyang, W., Mueller, F., Hjelmare, M., Lundberg, E., and Zimmer, C. (2019). ImJoy: an open-source computational platform for the deep learning era. arxiv.

Parker, R. and Sheth, U. (2007). P bodies and the control of mRNA translation and degradation. Molecular Cell, 25:(5)635–646.

Phillips, C. M., Montgomery, T. A., Breen, P. C., and Ruvkun, G. (2012). MUT-16 promotes formation of perinuclear Mutator foci required for RNA silencing in the *C. elegans* germline. Genes and Development, 26:(13)1433–1444.

Putker, M., Madl, T., Vos, H. R., de Ruiter, H., Visscher, M., van den Berg, M. C. W., Kaplan, M., Korswagen, H. C., Boelens, R., Vermeulen, M., Burgering, B. M. T., and Dansen, T. B. (2013). Redox-dependent control of FOXO/DAF-16 by transportin-1. Molecular Cell, 49:(4)730–742.

Raj, A. and Tyagi, S. (2010). Detection of individual endogenous RNA transcripts in situ using multiple singly labeled probes. Methods in enzymology, 472:(10)365–386.

Raj, A., van den Bogaard, P., Rifkin, S. A., van Oudenaarden, A., and Tyagi, S. (2008). Imaging individual mRNA molecules using multiple singly labeled probes. Nature Methods, 5:(10)877–879.

Robertson, S. and Lin, R. (2015). The Maternal-to-Zygotic Transition in *C. elegans*. In Current Topics in Developmental Biology, volume 113, pages 1–42. Academic Press Inc.

Samacoits, A., Chouaib, R., Safieddine, A., Traboulsi, A. M., Ouyang, W., Zimmer, C., Peter, M., Bertrand, E., Walter, T., and Mueller, F. (2018). A computational framework to study sub-cellular RNA localization. Nature Communications, 9(1).

Sawyer, J. M., Glass, S., Li, T., Shemer, G., White, N. D., Starostina, N. G., Kipreos, E. T., Jones, C. D., and Goldstein, B. (2011). Overcoming redundancy: An RNAi enhancer screen for morphogenesis genes in *Caenorhabditis elegans*. Genetics, 188:(3)549–564.

Schindelin, J., Arganda-Carreras, I., Frise, E., Kaynig, V., Longair, M., Pietzsch, T., Preibisch, S., Rueden, C., Saalfeld, S., Schmid, B., Tinevez, J. Y., White, D. J., Hartenstein, V., Eliceiri, K., Tomancak, P., and Cardona, A. (2012). Fiji: An open-source platform for biological-image analysis. Nature Methods, 9:(7)676–682.

Schisa, J. A., Pitt, J. N., and Priess, J. R. (2001). Analysis of RNA associated with P granules in germ cells of *C. elegans* adults. Development (Cambridge, England), 128:(8)1287–1298.

Schulz, K. N. and Harrison, M. M. (2019). Mechanisms regulating zygotic genome activation. Nature Reviews Genetics, 20:(4)221–234.

Seydoux, G. (2018). The P granules of *C. elegans*: A genetic model for the study of RNA–protein condensates. Journal of Molecular Biology, 430:(23)4702–4710.

Seydoux, G. and Fire, A. (1994). Soma-germline asymmetry in the distributions of embryonic RNAs in *Caenorhabditis elegans*. Development, 120:(10)2823–2834.

Seydoux, G., Mello, C. C., Pettitt, J., Wood, W. B., Priess, J. R., and Fire, A. (1996). Repression of gene expression in the embryonic germ lineage of *C. elegans*. Nature, 382:(6593)713–716.

Shaffer, S. M., Wu, M. T., Levesque, M. J., and Raj, A. (2013). Turbo FISH: A method for rapid single molecule RNA FISH. PLoS ONE, 8(9).

Sheth, U., Pitt, J., Dennis, S., and Priess, J. R. (2010). Perinuclear P granules are the principal sites of mRNA export in adult *C. elegans* germ cells. Development, 137:(8)1305–1314.

Shimada, M., Kawahara, H., and Doi, H. (2002). Novel family of CCCH-type zinc-finger proteins, MOE-1, −2 and −3, participates in *C. elegans* oocyte maturation. Genes to Cells, 7:(9)933–947.

Spike, C. A., Coetzee, D., Nishi, Y., Guven-Ozkan, T., Oldenbroek, M., Yamamoto, I., Lin, R., and Greenstein, D. (2014). Translational control of the oogenic program by components of OMA ribonucleoprotein particles in *Caenorhabditis elegans*. Genetics, 198:(4)1513–1533.

Strome, S. and Wood, W. B. (1982). Immunofluorescence visualization of germ-line-specific cytoplasmic granules in embryos, larvae, and adults of *Caenorhabditis elegans*. Proceedings of the National Academy of Sciences, 79:1558–1562.

Subramaniam, K. and Seydoux, G. (1999). *nos-1* and *nos-2*, two genes related to *Drosophila nanos*, regulate primordial germ cell development and survival in *Caenorhabditis elegans*. Development (Cambridge, England), 4871:(21)4861–4871.

Sweede, M., Ankem, G., Chutvirasakul, B., Azurmendi, H. F., Chbeir, S., Watkins, J., Helm, R. F., Finkielstein, C. V., and Capelluto, D. G. S. (2008). Structural and membrane binding properties of the prickle PET domain. Biochemistry, 47:(51)13524–13536.

Tenenhaus, C., Subramaniam, K., Dunn, M. A., and Seydoux, G. (2001). PIE-1 is a bifunctional protein that regulates maternal and zygotic gene expression in the embryonic germ line of *Caenorhabditis elegans*. Genes and Development, 15:(8)1031–1040.

Tintori, S. C., Osborne Nishimura, E., Golden, P., Lieb, J. D., and Goldstein, B. (2016). A transcriptional lineage of the early *C. elegans* embryo. Developmental Cell, 38:(4)430–444.

Trcek, T., Grosch, M., York, A., Shroff, H., Lionnet, T., and Lehmann, R. (2015). *Drosophila* germ granules are structured and contain homotypic mRNA clusters. Nature Communications, 6.

Tsanov, N., Samacoits, A., Chouaib, R., Traboulsi, A. M., Gostan, T., Weber, C., Zimmer, C., Zibara, K., Walter, T., Peter, M., Bertrand, E., and Mueller, F. (2016). SmiFISH and FISH-quant - A flexible single RNA detection approach with super-resolution capability. Nucleic Acids Research, 44(22).

Updike, D. L., Knutson, A. K. A., Egelhofer, T. A., Campbell, A. C., and Strome, S. (2014). Germ-granule components prevent somatic development in the *C. elegans* germline. Current Biology, 24:(9)970–975.

Van Fürden, D., Johnson, K., Segbert, C., and Bossinger, O. (2004). The *C. elegans* ezrin-radixin-moesin protein ERM-1 is necessary for apical junction remodelling and tubulogenesis in the intestine. Developmental Biology, 272:(1)262–276.

Vastenhouw, N. L., Cao, W. X., and Lipshitz, H. D. (2019). The maternal-to-zygotic transition revisited. Development, 146(11):dev161471.

Voronina, E., Paix, A., and Seydoux, G. (2012). The P granule component PGL-1 promotes the localization and silencing activity of the PUF protein FBF-2 in germline stem cells. Development (Cambridge), 139:(20)3732–3740.

Voronina, E., Seydoux, G., Sassone-Corsi, P., and Nagamori, I. (2011). RNA granules in germ cells. Cold Spring Harbor Perspectives in Biology, 3(12).

Walker, A. K., Boag, P. R., and Blackwell, T. K. (2007). Transcription reactivation steps stimulated by oocyte maturation in *C. elegans*. Developmental Biology, 304:(1)382–393.

Wan, G., Fields, B. D., Spracklin, G., Shukla, A., Phillips, C. M., and Kennedy, S. (2018). Spatiotemporal regulation of liquid-like condensates in epigenetic inheritance. nature, 557:(7707)679–683.

Wang, J. T., Smith, J., Chen, B. C., Schmidt, H., Rasoloson, D., Paix, A., Lambrus, B. G., Calidas, D., Betzig, E., and Seydoux, G. (2014). Regulation of RNA granule dynamics by phosphorylation of serine-rich, intrinsically disordered proteins in *C. elegans*. eLife, 3:1–23.

Zhang, F., Barboric, M., Blackwell, T. K., and Peterlin, B. M. (2003). A model of repression: CTD analogs and PIE-1 inhibit transcriptional elongation by P-TEFb. Genes and Development, 17:(6)748–758.

Zhang, Y., Foster, J. M., Kumar, S., Fougere, M., and Carlow, C. K. (2004). Cofactor-independent phosphoglycerate mutase has an essential role in *Caenorhabditis elegans* and is conserved in parasitic nematodes. Journal of Biological Chemistry, 279:(35)37185–37190.

Zhang, Y., Foster, J. M., Nelson, L. S., Ma, D., and Carlow, C. K. (2005). The chitin synthase genes *chs-1* and *chs-2* are essential for *C. elegans* development and responsible for chitin deposition in the eggshell and pharynx, respectively. Developmental Biology, 285:(2)330–339.

