## SupplementalFigures for "mRNA localization is linked to translation regulation in the *Caenorhabditis elegans* germ lineage"

### Supplement

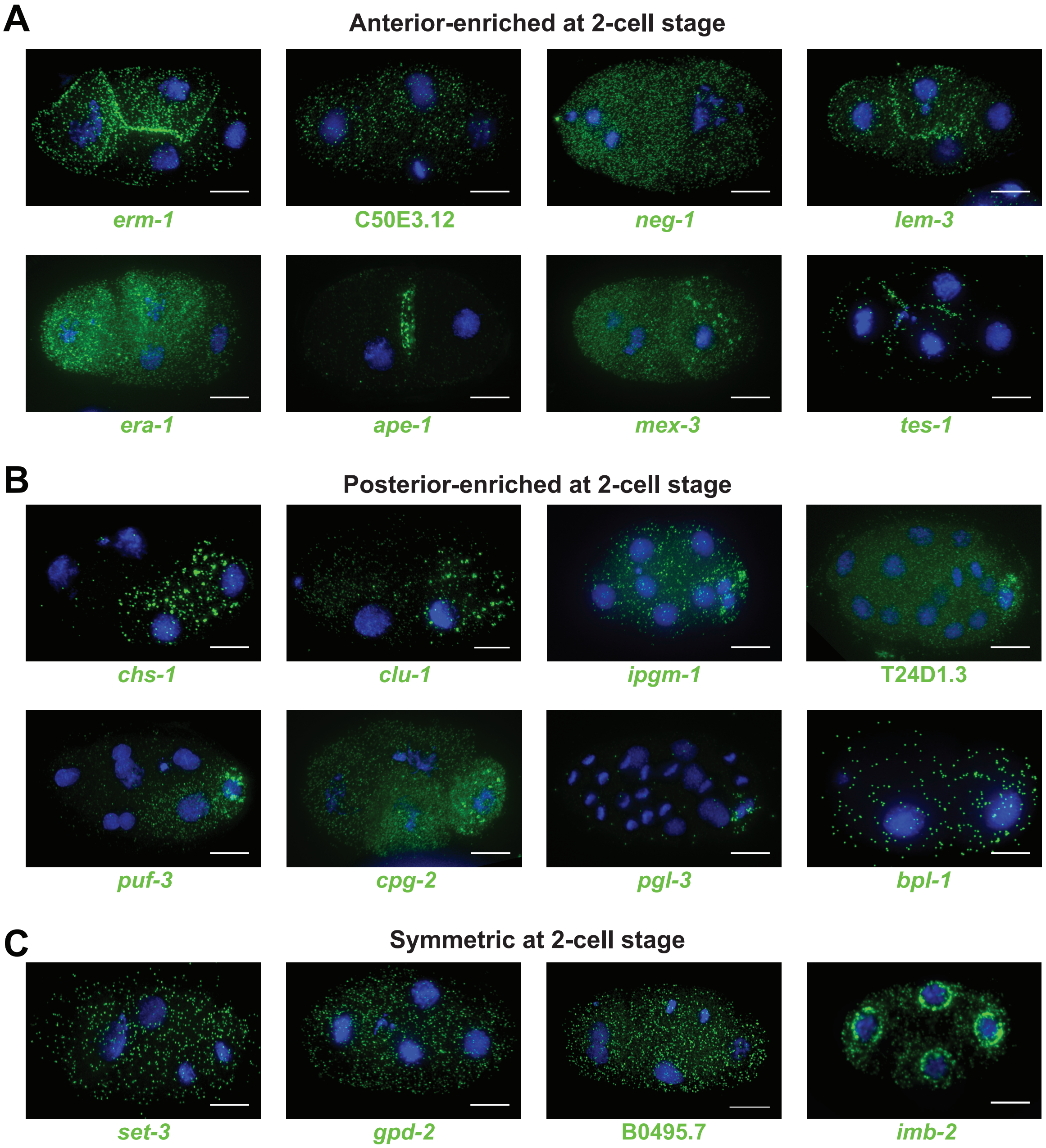


**Figure S1. Subcellular localization patterns of surveyed mRNA transcripts.** 8 AB-enriched, 8 $P_1$ enriched, and 4 symmetric transcripts were selected examination by smFISH with all transcripts in green and DAPI in blue. **(A)** AB-enriched transcripts *erm-1 (Ezrin/Radixi/Moesin)*, C50E3.12, *neg-1 (Negative Effect on Gut development, lem-3 (LEM domain protein), era-1 (Embryonic mRna Anterior), ape-1 (APoptosis Enhancer), mex-3 (Muscle EXcess)*, and *tes-1(TEStin homolog)* are shown. **(B)** $P_1$-enriched transcripts *chs-1 (CHitin Synthase), clu-1 (yeast CLU related), ipgm-1 (cofactor Independent PhosphoGlycerate Mutase homolog)*, T24D1.3, *puf-3 (PUF domain containing), cpg-2 (Chondroitin ProteoGlycan), pgl-3 (P-GranuLe abnormality), and bpl-1 (Biotin Protein Ligase)* are shown. **(C)** Uniformly distributed transcripts *set-3 (SET domain containing), gpd-2 (Glyceraldehyde 3-Phosphate Dehydrogenase)*, B0495.7, *imb-2 (IMportin Beta family)* are shown. Tabulation of the results are in Table 1.

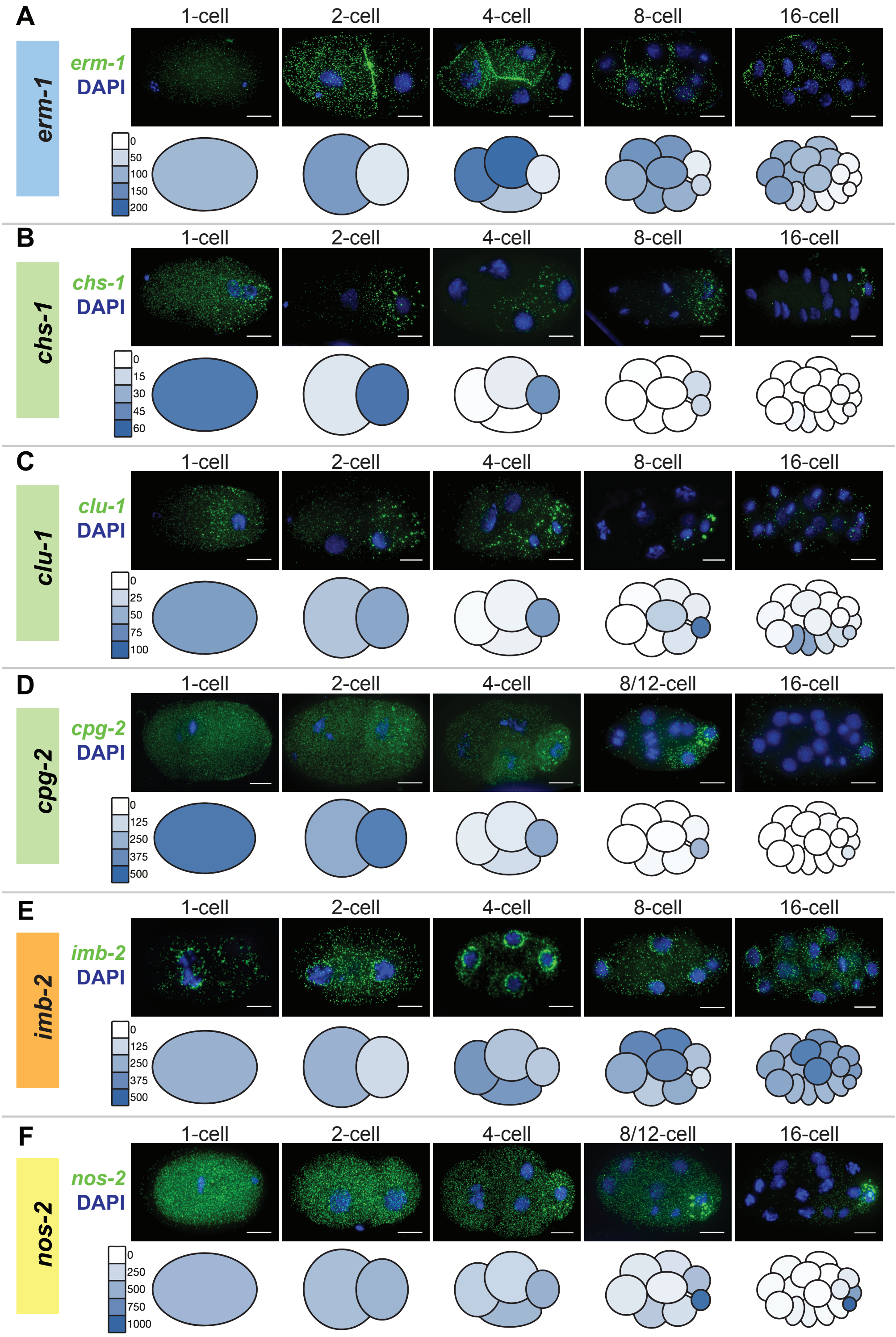


**Figure 2. Localization patterns of transcripts over developmental time.** smFISH microscopy localizations of *erm-1* **(A)**, *chs-1* **(B)**, *clu-1* **(C)**, *cpg-2* **(D)**, *imb-2* **(E)**, and *nos-2* **(F)** shown from 1-cell stage zygotes to the 16-stage. mRNA signal is in green. DAPI is in blue. Below each series of images is single-cell RNA-seq data for the same transcript [Tintori et al., 2016].

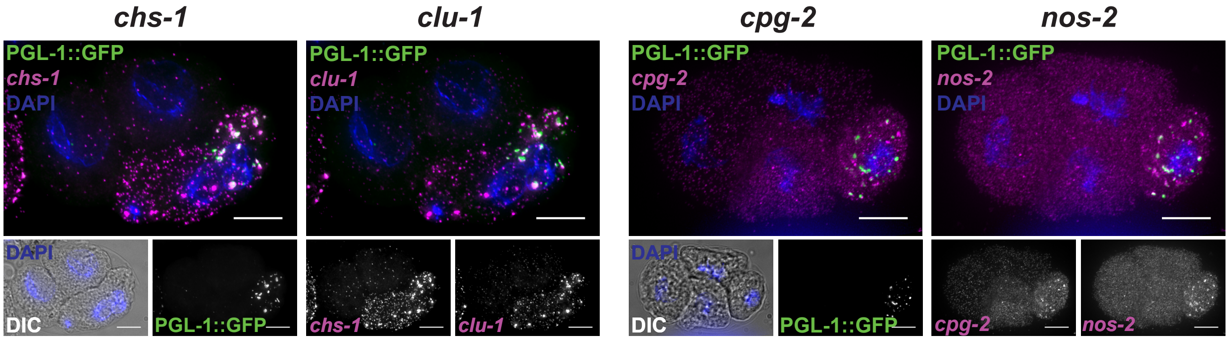


**Figure 3. $P_1$-enriched, clustered transcripts co-localize with the P granule marker PGL-1::GFP.** In addition to co-localizing with GFP signal in the P granule marker strain containing GLH-1::GFP (Figure 3), *chs-1, clu-1, cpg-2,* and *nos-2* mRNAs (all in magenta) also co-localize with a second P granule marker protein, PGL-1::GFP (green). DAPI is illustrated in blue to visualize nuclei and illustrate the 4-cell stage of development.

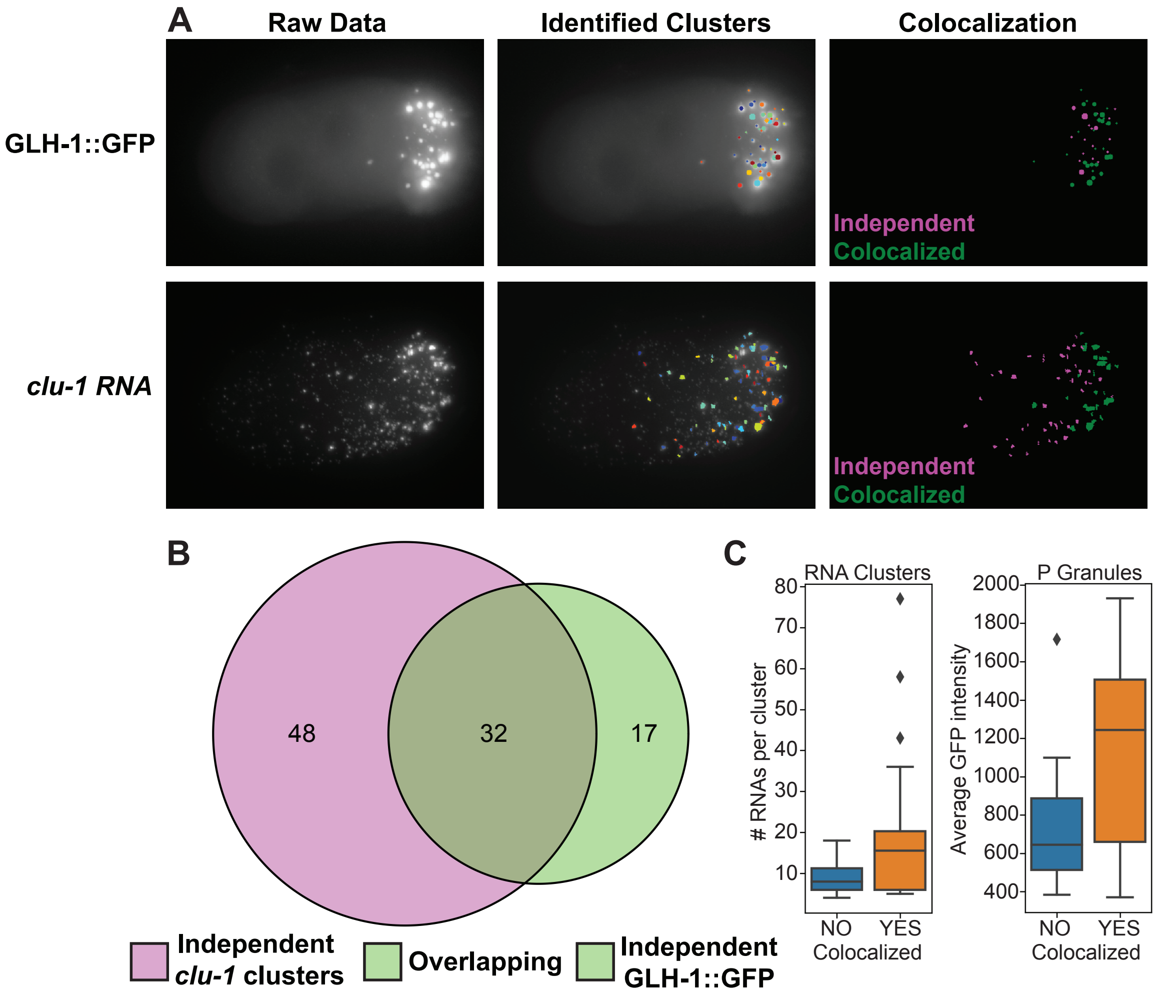


**Figure 4. Quantification of mRNA cluster overlap with the P granules.** mRNA cluster overlap with GLH-1::GFP labeled P granules is calculated using micrographs of GLH-1::GFP and clustered RNAs (*clu-1* shown) **(A, left)**, computationally identifying P granules and RNA clusters **(A, middle)**, and comparing the 3D masks for overlap to identify independent P granules and RNA clusters (magenta) or colocalized clusters (green) **(A, right)**. **(B)** A Venn-Euler diagram illustrating the number of independent *clu-1* mRNA clusters (magenta), independent P granules (light green), and overlapping P granules and mRNA clusters (dark green) in a single embryo (from A). **(C)** Box plots comparing the size of non-overlapping mRNA clusters and P granules to those overlapping shows larger mRNA clusters more commonly overlap with P granules (left) and brighter P granules more commonly overlap with mRNA clusters (right).

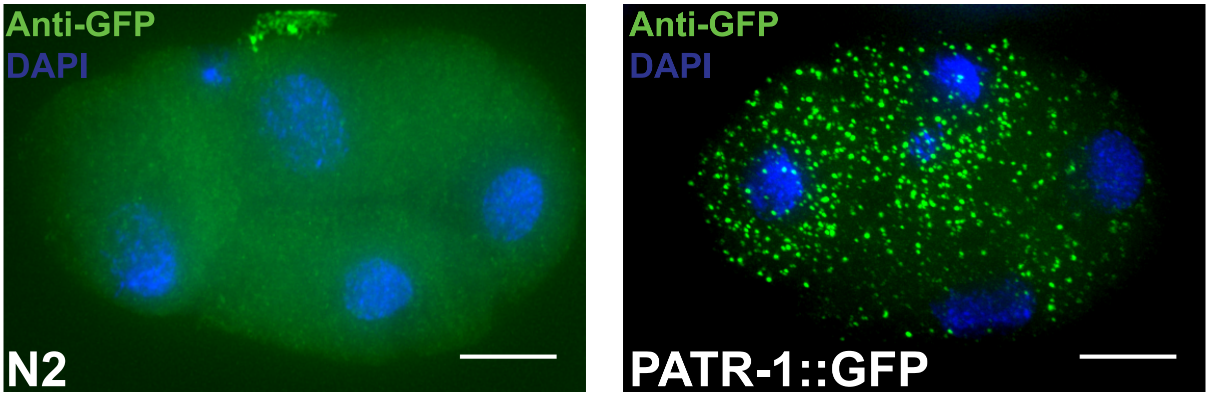


**Figure 5. Immunofluorescence control images for anti-GFP antibody staining.** The anti-GFP antibody reports PATR-1::GFP localization (green, right) in PATR-1::GFP containing strains as compared to N2 wild type un-transformed control strains (green, left). DAPI staining is shown in blue to visualize nuclei and illustrate the 4-cell stage of development.

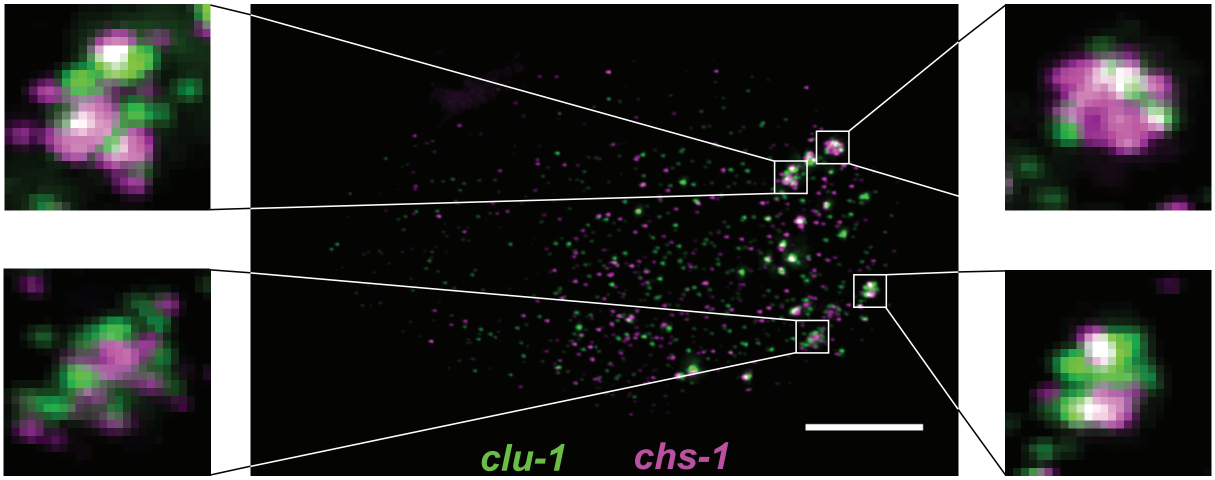


**Figure 6. mRNA clusters display homotypic clustering within P granules.** *chs-1* mRNA (magenta) tend to homotypically cluster in the core of P granules while *clu-1* mRNA (green) also cluster homotypically, but near the peripheries of P granules.

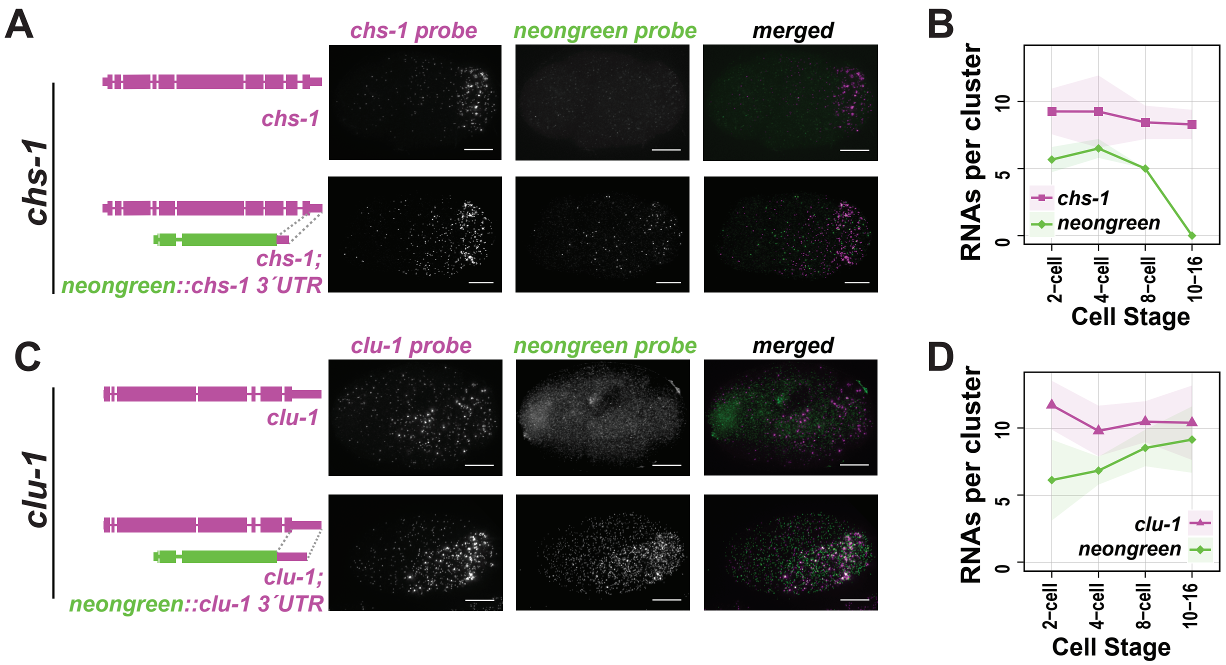


**Figure 7.** ***chs-1* and *clu-1* 3'UTRs direct neongreen mRNAs to subcellular clusters.** **(A)** A single copy of the *chs-1* 3'UTR appended to *monomeric NeonGreen (mex-5p::mNeonGreen::chs-1 3'UTR)* was transgenically introduced into otherwise wild-type worms to test whether the 3'UTR of *chs-1* was sufficient to drive the cell peripheral localization pattern observed for full length *chs-1* mRNA. Wild-type (above) and transgenic (below) 4-cell stage embryos were imaged by smFISH using probes to the endogenous *chs-1* mRNA (left), *mNeonGreen* mRNA (middle), and the channels were then merged (right). **(B)** Quantification of (A) over developmental time. **(C)** As in (A) with using the *clu-1* 3'UTR sequence. **(D)** As in (B) using the *clu-1* 3'UTR.

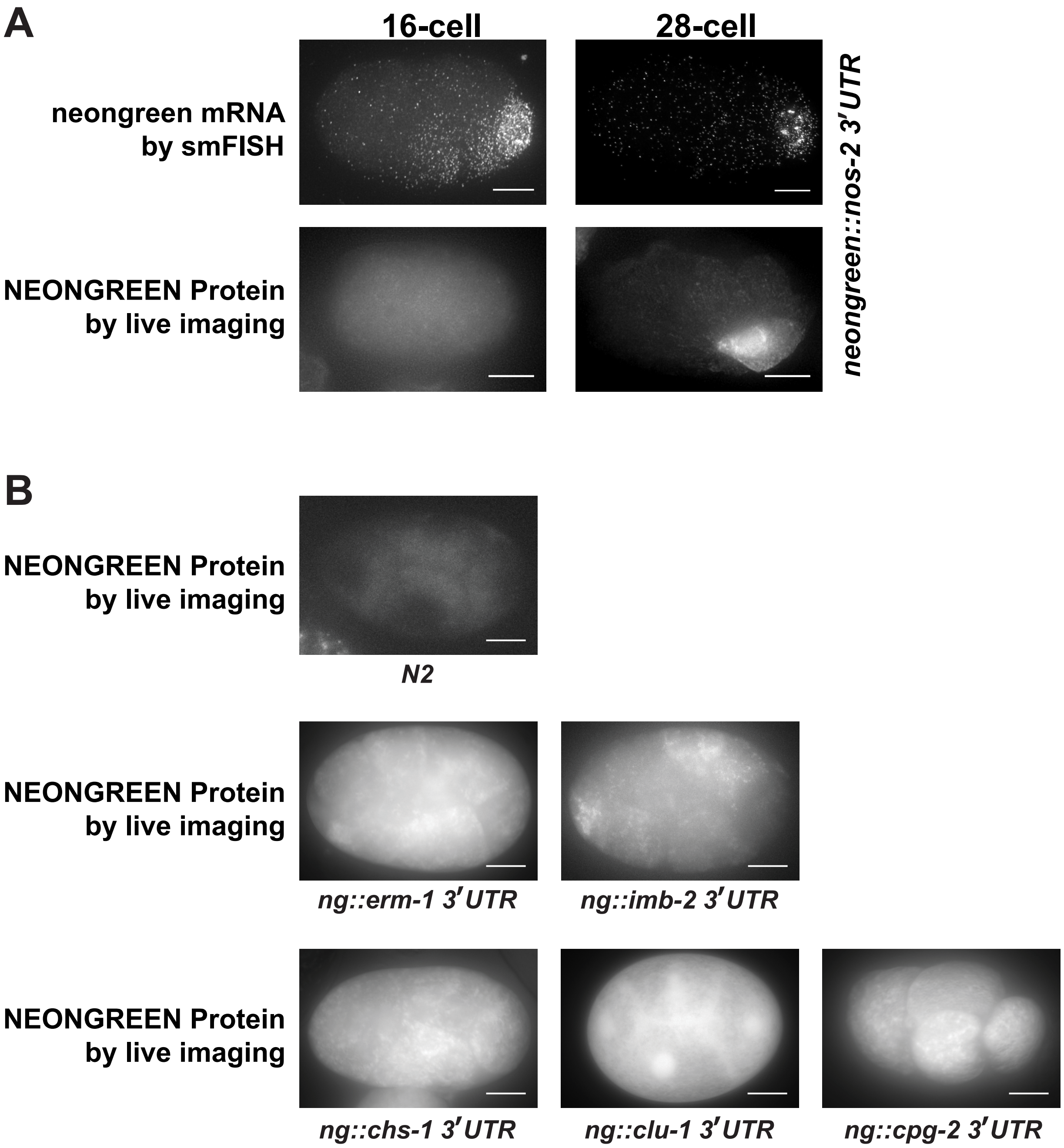


**Figure 8.** ***Pmex-5::mNeonGreen::nos-2 3'UTR* RNA recapitulates endogenous translation repression and activation.** **(A, left)** A *Pmex-5::mNeonGreen::nos-2 3'UTR* embryo at the 16-cell stage. smFISH for *mNeonGreen* RNA demonstrated normal RNA localization. Epifluorescent microscopy of living *Pmex-5::mNeonGreen::nos-2 3'UTR* embryos at the 16-cell stage showed no expression of the mNeonGreen reporter protein. **(A, right)** As in (A) at the 28-cell stage. Epifluorescent microscopy of living *Pmex-5::mNeonGreen::nos-2 3'UTR* embryos at the 28-cell stage showed P lineage specific expression of the mNeonGreen reporter protein. **(B)** Epifluorescent microscopy of wildtype N2 embryos demonstrates no apparent fluorescence while *Pmex-5::mNeonGreen::3'UTR of Interest* embryos show low levels of cell-non-specific mNeonGreen fluorescence.

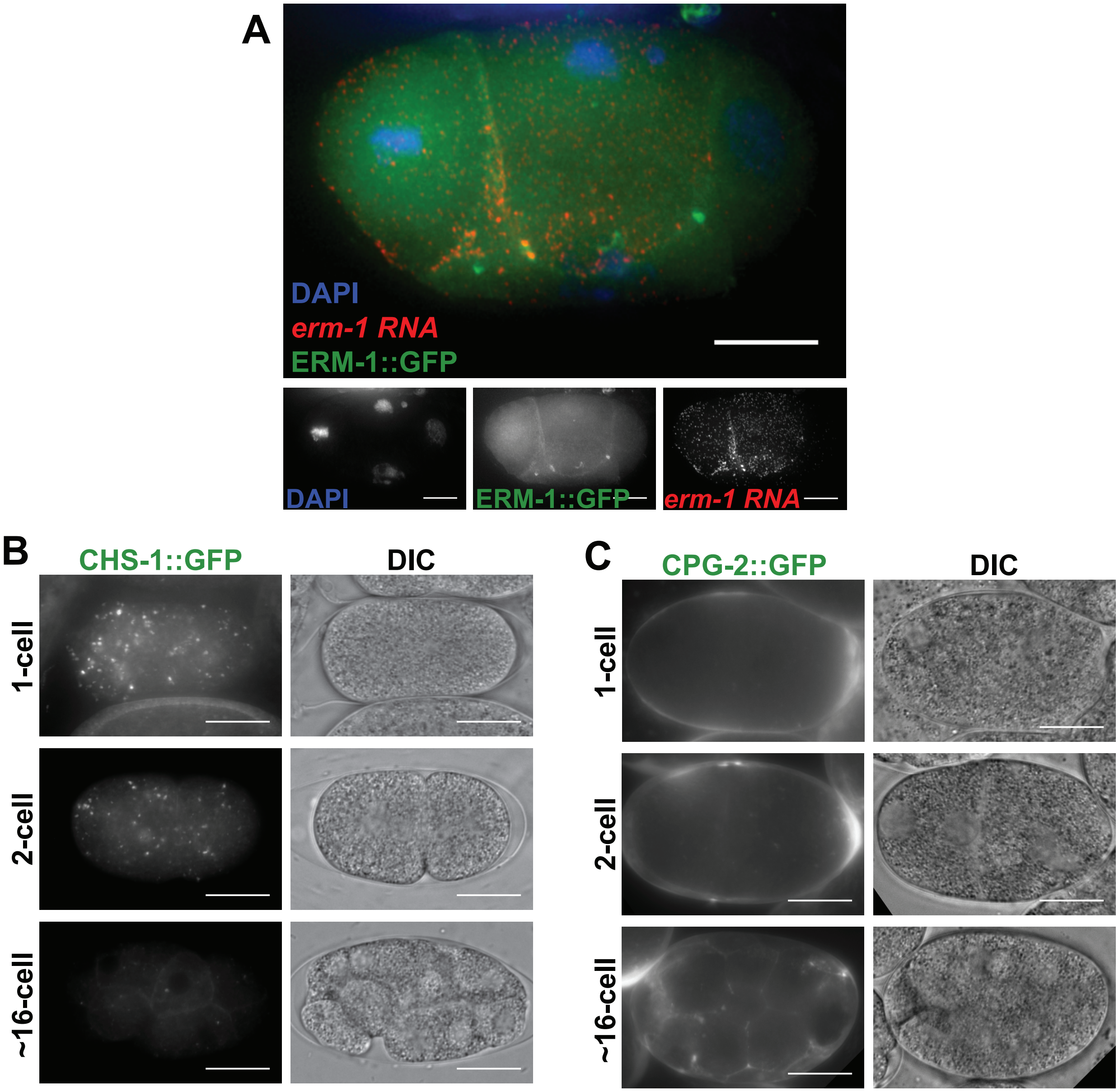


**Figure 9.** ***erm-1::GFP* RNA localizes like endogenous *erm-1*.** **(A)** smFISH microscopy of *Perm-1::erm-1 ORF::GFP::erm-1 3'UTR* RNA (green) colocalizes with endogenous *erm-1* RNA (red). **(B)** Epifluorescent microscopy of CHS-1::GFP embryos at the 1, 2, and 16-cell stages of embryogenesis show gradual depletion of the CHS-1::GFP protein puncta. **(C)** As in (B) imaging CPG-2::GFP embryos. CPG-2::GFP protein can be seen in the extracellular space of the embryo, but not within cells.

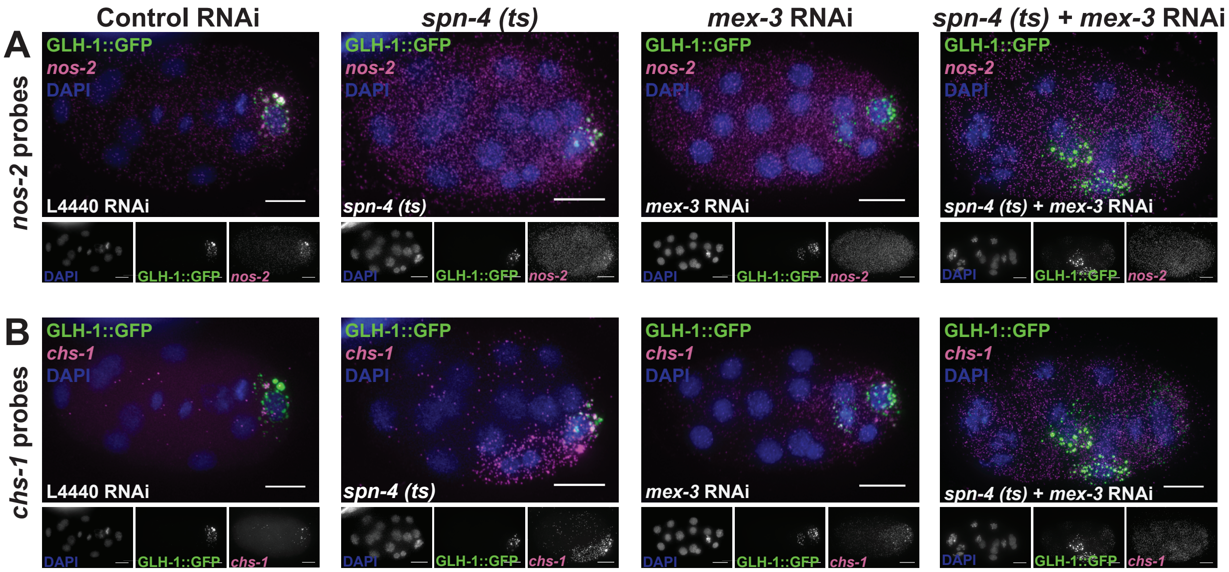


**Figure 10. Knockdown of *spn-4* and *mex-3* simultaneously resulted in embryonic defects and mislocalization of RNA.** GLH-1::GFP and GLH-1::GFP *spn-4(or191)* temperature sensitive embryos raised at 25°C (GLH-1::GFP shown in green) were treated with L4440 or *mex-3* RNAi and probed for *chs-1* **(A)** and *nos-2* **(B)** mRNAs by smFISH (magenta). DNA was also stained with DAPI (blue). GLH-1::GFP *spn-4(or191)* embryos raised at 15°C and treated with either L4440 or *mex-3* RNAi phenocopied GLH-1::GFP embryos raised at 25°C with the same RNAi treatment (data not shown).

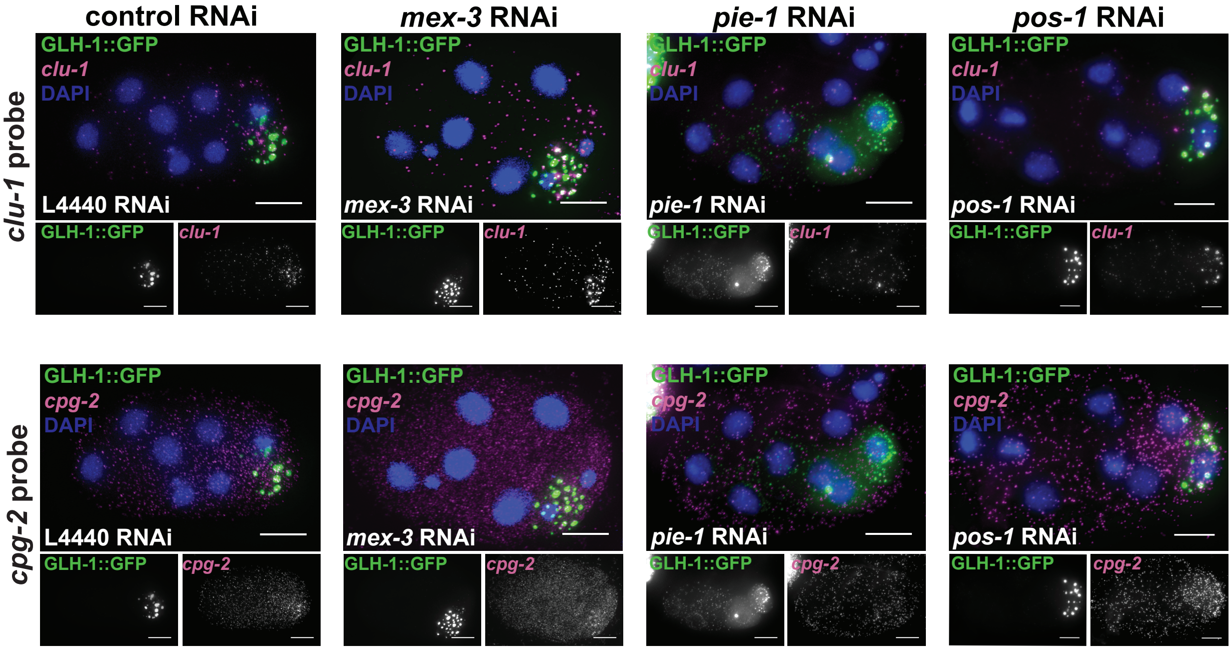


**Figure 11. Knockdown of RBPs important for *nos-2* transcript localization results in perturbed *cpg-2* and *clu-1* transcript localization. (A)** GLH-1::GFP (green) embryos were treated with L4440, *mex-3, pie-1,* or *pos-1* RNAi and probed for *clu-1* **(A)** and *cpg-2* **(B)** mRNAs by smFISH (magenta). DNA was also stained with DAPI (blue).

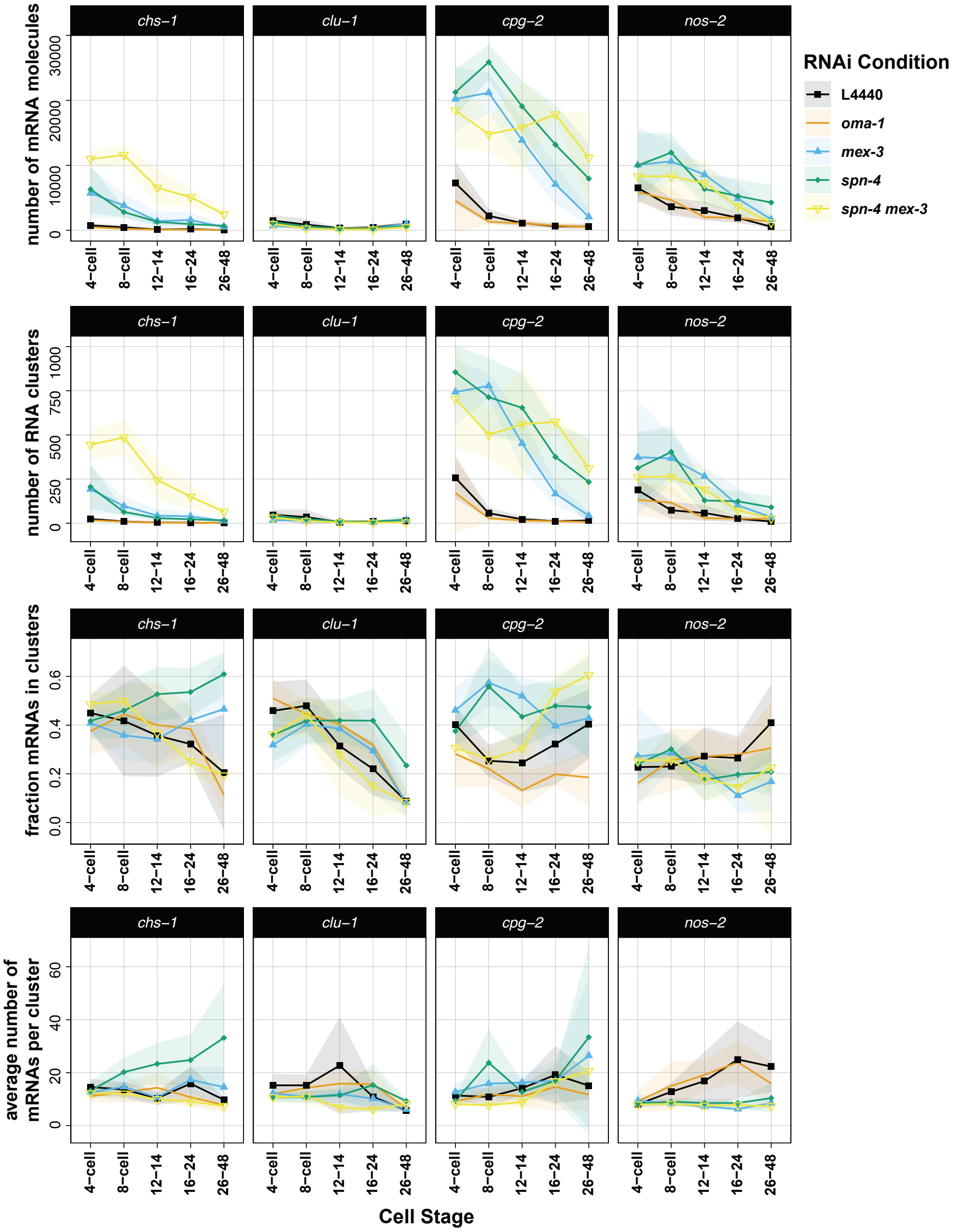


**Figure 12. Quantification of mRNA clustering under RBP knockdown conditions** Statistical metrics characterizing mRNA clustering are shown: 1) the total number of RNAs in each embryo, 2) the total number of clusters identified in each embryo, 3) the fraction of total mRNAs located within clusters (as opposed to cluster-independent), and 4) the average estimated number of mRNA molecules per cluster within a given embryo. Four transcripts were assayed in five knockdown conditions over 5 developmental time points. The average of each metric (line) and their standard deviation (shading) are shown representing a minimum of 5 embryos assayed.
